# Molecular requirements of chromogranin B for the long-sought anion shunter of regulated secretion

**DOI:** 10.1101/2024.12.24.630220

**Authors:** Gaya P. Yadav, Mani Annamalai, D. Walker Hagan, Lina Cui, Clayton Mathews, Qiu-Xing Jiang

## Abstract

All eukaryotes utilize regulated secretion to release molecular signals packaged in secretory granules for local and remote signaling. An anion shunt conductance was first suggested in secretory granules of bovine chromaffin cells nearly five decades ago. Biochemical identity of this conductance remains undefined. CLC-3, an intracellular Cl^-^/H^+^ exchanger, was proposed as a candidate sixteen years ago, which, however, was contested experimentally. Here, we show that chromogranin B (CHGB) makes the kernel of the long-sought anion shunter in cultured and primary neuroendocrine cells and its channel functions are essential to proper granule maturation. Intragranular pH measurements and cargo maturation assays revealed that normal granular acidification, proinsulin-insulin conversion, and dopamine-loading in neuroendocrine cells all rely on functional CHGB+ channels. Primary β-cells from *Chgb-/-* mice exhibited persistent granule deacidification, which suffices to uplift plasma proinsulin level, diminish glucose-induced 2^nd^-phase insulin secretion and dwindle monoamine content in chromaffin granules from the knockout mice. Data from targeted genetic manipulations, dominant negativity of a deletion mutant lacking channel-forming parts and tests of CLC-3/5 and ANO-1/2 all exclude CHGB*-less* channels from anion shunting in secretory granules. The highly conserved CHGB+ channels thus function in regulated secretory pathways in neuronal, endocrine, exocrine and stem cells of probably all vertebrates.

**HIGHLIGHTS:**

- Loss of CHGB channel functions impairs secretory granule acidification in neuroendocrine cells, which necessitates anion shunt conduction.
- CHGBΔMIF, a mutant unable to form a functional Cl^-^ channel, exerts negative dominance on endogenous CHGB and results in granule deacidification in cultured cells.
- Neither CLC-3 & -5 nor ANO-1 & -2 participate in the CHGB-mediated granule acidification. Clcn3 knockout effects on regulated secretion can be attributed to its functions in endosomal and endolysosomal compartments.
- Primary *Chgb-/-* β-cells exhibit persistent granule deacidification, presenting a unifying mechanism for disparate mouse phenotypes: hyperproinsulinemia, near abrogation of 2^nd^ phase insulin release after glucose challenge and diminution of monoamine contents in chromaffin granules.

## INTRODUCTION

Eukaryotic organisms have two general types of secretory pathways: constitutive and regulated secretion. [1] All human tissues and organs contain regulated secretory cells, such as neurons and neuroendocrine cells.[2-6] Regulated secretion releases molecular signals that act at both local and systems levels to achieve physiological homeostasis.[7] These signals, including proteins, short peptides, ions, small organic compounds, etc., are stored in membrane-enclosed vesicles. Three well-known types of regulated secretory vesicles are synaptic vesicles in neurons, lysosome-related organelles (LRO) in immune granulocytes, and secretory granules (SGs) in exocrine, endocrine, neuronal and stem cells.[8-12] Among these three, the SGs are more generic because their secretion was first employed to unveil intracellular vesicle trafficking from ER to Golgi to plasma membranes by Palade and his colleagues and is conserved among all eukaryotes.[11, 13-17] Regulated secretion includes three intracellular steps: biogenesis of immature secretory granules (ISGs) at the *trans*-Golgi network (TGN), maturation of ISGs into dense-core SGs (DCSGs), and stimulus-elicited exocytosis of DCSGs.[1, 18] Even though protein machines for vesicular trafficking have been studied for decades,[19-21] molecular mechanisms underlying the three steps of regulated secretion remain inadequately characterized.

Anion shunt conduction across granular membranes is essential to SG maturation with interior acidity driven by vacuolar H^+^-ATPase (vATPase).[22] For each 100-nm DCSG with an estimated interior protein concentration of ∼150 mg/ml, vATPases need to translocate 10^5^ – 10^6^ protons to acidify the granular lumen from pH ∼6.5 to ∼5.5,[22, 23] whereas merely 200 positive elementary charges are sufficient to establish a positive potential of ∼100 mV that is strong enough to deter H^+^ pumping completely. The surplus positive charge must hence be neutralized by ion flow, either cation efflux or anion influx or both, for acidification to happen. A Cl^-^ influx into SGs was first discovered in late 1970s in bovine chromaffin granules as a shunt current to nullify charge accumulation.[22, 24-26] Besides electrochemical effects, the acidic milieu in SGs catalyzes cargo maturation via proteolysis or molecular transport across membranes and compaction of granular contents, via protein salting-out, liquid-liquid phase separation (LLPS), cation binding or amyloidosis. [27-30] The transmembrane proton gradient supports energetically translocation of small organic molecules and/or divalent cations from cytosol into SGs. Normal maturation of secretory granules and processing of cargo molecules are therefore tightly coupled to granule acidification. Without anion shunt currents, H^+^- pumping would slow down quickly until being fully abolished because a positive potential of +60 - 100 mV would develop immediately with charge accumulated.[31, 32] The anion shunt conduction is thus of critical importance to regulated secretion.

Biochemical basis of the anion shunt conductance has been mysterious for nearly 50 years. CLC-3, which was first thought of as a Cl^-^ channel, was proposed as a candidate, but its physical presence in secretory granules was contested due to antibody nonspecificity.[33-37] Further, instead of being a channel, CLC-3 was later identified as a Cl^-^/H^+^ (2:1) exchanger with a fS conductance, making it unfit for fast charge neutralization in a SG unless its abundance could be three to four orders of magnitude higher to match a channel conductance.[38, 39] More permeation of organic anions, such as SCN^-^, over Cl^-^ through CLC-3 is also inconsistent with the need to avoid enriching organic anions into SGs.[40] On the other hand, prior studies found little cation permeation (except H^+^) in chromaffin granules,[25] suggesting that certain cation channels indicated by analysis of electrical recordings from isolated granules may not be abundant enough for ion shunting, either.[41-45] Furthermore, potential contamination of purified SGs by other membranes[46] and varied antibody specificity introduced more uncertainty in the candidate approaches,[35, 47-49] leaving a significant gap in biochemical identity of the anion shunt conductance.

Chromogranin B (CHGB, also abbreviated as CgB in the past) as an alternative candidate for the anion shunt conductance still requires direct physiological tests.[50] By serendipity, we found that CHGB has soluble and membrane-integral forms, the latter alone reconstitutes Cl^-^ channels in proteoliposomes and planar bilayers and the CHGB-containing (CHGB+) channels are active on the cell surface after granule exocytosis.[51, 52] Such a membrane-resident channel suggests an unusual transmembrane topology, which, like other dimorphic proteins, is stabilized by hydrophobic or amphipathic segments.[53] The CHGB transmembrane topology differs surely from surface interactions with lipids or membrane proteins proposed by two groups in early 1990s,[54-56] to agree with a presumed soluble state (as oligomers) of CHGB.[57-59] As an intrinsic property, membrane integration of CHGB can account for the anion channel activities, nearly half protein at or near granular membrane in immuno-EM studies, CHGB puncta on the cell surface after granule release and whole-cell recordings of large anionic currents. All these results point to an appealing proposition that CHGB might be an integral part of the anion shunt conductance, which, however, still face questions on whether special membrane composition and/or luminal constituents of SGs might suppress the CHGB+ channel functions *in situ*. Direct tests are hence desired but are still missing.

In the present study, we built up on our previous work to investigate the physiological roles of the proposed CHGB+ channels in SGs. We studied the effects of losing CHGB or its channel functions based on the tight coupling among anion shunting, granule acidification, cargo maturation or granule filling in neuroendocrine cells. We analyzed dominant negative effects of a deletion mutant on granule acidification and assessed impairment of granule acidification in primary neuroendocrine cells from *Chgb*-null mice. All our data converge to a unifying hypothesis that CHGB constitutes the kernel of the anion shunt conductance in SGs, which not only resolves a long-standing mystery in neurons and neuroendocrine cells and in cell biology at large, but also unifies seemingly discordant phenotypes of *Chgb-/-* mice.[60-62]

## RESULTS

### CHGB knockdown impairs granule acidification and proinsulin-to-insulin conversion

Our first unknown is whether the CHGB+ channels detectable on the cell surface after granule exocytosis are active in SGs inside cells or not (Fig. 1A). The tight coupling of granule acidification and cargo maturation to anion shunt conduction makes it feasible to assess the physiological effects of losing the channel functions despite that SGs in intact cells are too small and inaccessible to direct electric recordings. We utilized insulin- and dopamine-secreting cells as model systems because they represent two typical types of regulated secretory pathways releasing small proteins (or peptides) and organic compounds, respectively. We employed the same siRNAs verified previously to knock down CHGB and tried to rescue by overexpressing wild-type and mutant forms (Fig. 1B).[51]

**Figure 1.**
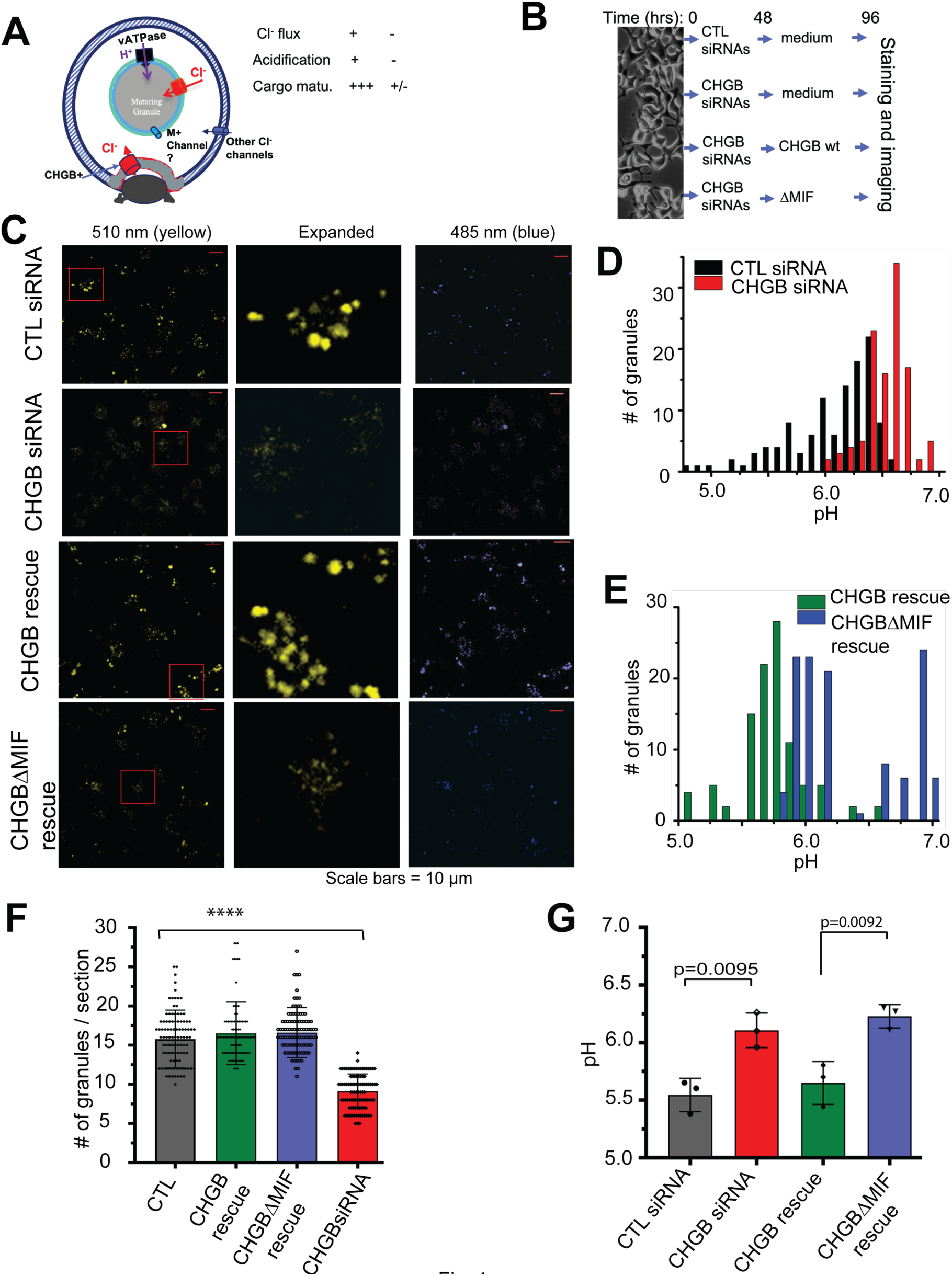
CHGB for normal granule acidification and insulin maturation in INS-1 cells. (**A**). A schematic diagram showing vATPase and the Cl^-^ shunt conductance in granular membranes, and the CHGB+ Cl^-^ channels delivered to the cell surface after granule release. The Cl^-^ shunt flux is coupled to granule acidification and cargo maturation. (**B**). Four different ways to treat cells before DND-160 staining for ratiometric pH measurements. Cells were transfected with control (CTL; top row) or CHGB-specific siRNAs (rows 2 to 4). In top two rows, cells were fed with fresh media after 48 hours and were stained after 96 hours. In rows 3 & 4, forty-eight hours after CHGB knockdown, the cells were transfected to overexpress CHGB or CHGBΔMIF and were imaged after another 48 hours. (**C**). Typical data of INS-1 cells treated as outlined in (**B**). Excitation: 410 nm; emission: 485 nm & 510 nm. A small area (red window) in each 510 nm image was expanded to show granules (the middle column). Measurement at 510-nm in rows 2 & 4 was weaker due to pH change. (**D**). Histograms of intragranular pH values from cells treated with CTL or CHGB siRNAs (black vs. red). (**E**). Histograms of intragranular pH values from CHGB knockdown cells overexpressing WT CHGB or CHGBΔMIF (green vs. blue bars), respectively. (**F**). Average numbers of granules per cell focused on a central section of cells treated as **B.** Errors: *s.d*. n ∼ 150. (**G**). Average pH values of secretory granules from cells as in **C-E** from 4 repeated experiments. Error: *s.d.,* n=4. **: *p* < 0.05; ***: *p* < 0.001; ****: *p* < 0.0001; *ns*: not significant.

Anion flux in SGs is coupled to granule acidification when proton-pumping works normally. It predicts that failure in proper granule acidification, called deacidification, reflects a significant loss of anion channel functions, which should impact on cargo maturation (Fig. 1A inset). Both SG acidification and cargo maturation can be (semi-)quantitatively measured. To monitor intragranular pH, we utilized DND-160, a ratiometric sensor whose variations in local loading do not affect pH readouts.[63] As tests of suitability, we conducted a set of control experiments. Firstly, syncollin, a granular protein by default, was fused with a fluorescent protein, pHluorin[64], and transiently expressed to label SGs before DND-160 staining (Supplementary Fig. 1A). Confocal imaging data showed that syncollin and DND-160 overlapped well in dye-stained compartments, meaning that DND-160 stains the SGs well in the INS-1 cells probably because SGs in these cells outnumber other acidic compartments like lysosomes by 2-3 orders of magnitude.[63] Secondly, we counted DND-160-stained SGs per cell section in INS-1 and PC-12 cells treated with control and CHGB-targeting siRNAs and imaged by confocal fluorescence microscopy (Supplementary Figs. 1B-C). Compared to control cells, CHGB knockdown decreased the average SG number by ∼65% in INS-1 and ∼50% in PC-12 cells, respectively, consistent with CHGB’s partial, but nonessential contribution to granule biogenesis and the potential compensatory effects in *Chgb-/-* and *Chga/Chgb* double knockout mice.[60-62, 65, 66] Thirdly, as a test on granule biogenesis, INS-1 cells treated with 5.0 mM NH_4_Cl showed an average number of granules per cell section (∼16), close to that (∼18) in cells treated with control siRNA (Supplementary Figs. 1B vs. Fig. 1F). Fourth, when INS-1 cells stained with DND-160 for 5 and 30 minutes were compared, the granular pH distributions from the two sets of cells were the same, in accord with the expected property of a ratiometric dye. We thus used 5-minute staining for all following experiments. Fifth, as a positive control for granular pH readings, dual excitation experiments in INS-1 cells treated with NH_4_Cl (Supplementary Figs. 2A-C vs. Figs. 1C, 1D & 1E) were performed. To enhance consistency among different experiments and minimize systematic errors, we calibrated DND-160 readout curves in a pH range of 4.7 – 7.5 in every microscopic system (as an example, Supplementary Fig. 2D). Because of expected SG pH variation in live cells, measurements from hundreds of SGs out of dozens of cells in each experiment were plotted in a histogram (Supplementary Fig. 2B) to obtain a mean granular pH. The readouts from repeated experiments were then used for statistical comparison (Supplementary Fig. 2C). Such an analysis is equivalent to pooling together data from repeated experiments as random statistical samplings and is mathematically robust based on the large number theorem (see Methods). As expected, NH_4_Cl treatment increased the mean granular pH to ∼6.6 because NH_4_^+^ buffered SG pH near neutrality. Sixth, as another negative control, mean granular pH of control siRNA-treated cells was ∼5.5, close to normal cells (Supplementary Figs. 2A, 2C). These data demonstrated that DND-160 was suitable and reliable for measuring intragranular pH in our own experiments.

Next, we compared CHGB knockdown cells with those treated with control siRNAs using a dual-excitation protocol and found an increment of the average granular pH by ∼0.6 units (Supplementary Figs. 2A-C). Loss of CHGB thus caused significant granule deacidification. To minimize cytotoxicity of near-UV beam, we switched to a dual-emission protocol for DND-160 (rows 1 and 2 in Fig. 1C) and detected the same change in granular pH (∼0.7 pH units) between CHGB knockdown (<PH> ∼6.2, red in Fig. 1D) and control (<PH> ∼5.5, black in Fig. 1D) cells (statistical comparison in Fig. 1G). These data showed that the two protocols produced essentially the same results, demonstrating high reproducibility in granular pH quantification.

The granule deacidification induced CHGB knockdown may result from downregulated H^+^-pumping (vATPase) or loss of ion shunt currents, or both [22, 24]. Because in CHGB-knockdown cells vATPase was delivered to cell surfaces reliably upon stimulated granule release [51] and its H^+^-pumping activities in different organelles other than SGs do not require CHGB, it is unlikely that the granular deacidification caused by CHGB loss stemmed from defective vATPase *per se*. But instead, it was the loss of anion shunt currents that probably led to positive charge accumulation and granule deacidification (Fig. 1A) [22]. If so, this phenotype should be rescuable by over-expressed wild-type CHGB as observed in Figs. 1C, 1E & 1G.

The observed reduction in the average SG number in CHGB-knockdown cells (Supplementary Figs. 1B-C and Fig. 1F) led us to examine the feasibility to separate CHGB’s roles in channel formation from those in granule biogenesis. To do so, we took advantage of a nonconducting deletion mutant, murine (m) CHGBΔMIF, which lacks a critical membrane-insertion fragment (MIF).[52] We thus expected that mCHGBΔMIF behave differently from the wildtype in rescuing the CHGB knockdown cells (Figs. 1B & C).[52, 66] Satisfyingly, overexpression of either mCHGB or mCHGBΔMIF was sufficient to restore the average number of secretory granules per cell section (Figs. 1C & 1F), because CHGB and CHGBΔMIF both contain intact the N-terminal sorting signal for granule biogenesis and were expressed well to overwhelm the leftover siRNAs.[66] To assess the capacity of these two constructs in granule biogenesis without CHGB knockdown, we compared their induction of secretory granule-like vesicles (SGLVs) in fibroblast cells. As a DND-160 substitute we introduced ClopHensor, a fluorescent protein-based sensor for ratiometric pH measurements.[67] Overexpression of CHGB, CHGBΔMIF or their fusion proteins induced DND-160-stained large SGLVs in both cell lines (Supplementary Figs. 3A, 3B), which are much brighter than those in control cells. It is also interesting to note the SGLVs in fibroblasts were acidic to retain DND-160, although to a lesser extent than regular SGs in INS-1 cells (rows 3 and 4 in Supplementary Fig. 3A), probably because CHGB alone failed to recruit enough vATPase and other granular components into SGLVs in fibroblasts. These results further demonstrate that both WT CHGB and CHGBΔMIF are competent in SG biogenesis, which enables them to rescue granule biogenesis in CHGB knockdown cells (Figs. 1C, 1F).

CHGBΔMIF and WT CHGB behaved the opposite in rescuing the CHGB knockdown-induced granule deacidification (rows 3 & 4 in Fig. 1C, and Figs. 1E & 1G). WT CHGB restored granular <PH> to ∼5.6 (red vs. green in Fig. 1G). In contrast, CHGBΔMIF failed to do so, but instead worsened deacidification by increasing granular <PH> slightly higher than that of the knockdown cells (pH 6.3 vs. 6.1), indicating potentially a dominant negative effect (blue vs. red in Fig. 1G). If loss of CHGB suppresses anion shunt currents, CHGB either serves to the anion conductance itself or facilitates the delivery to SGs of a different anion conductance, whose conduction needs no CHGB. The complete rescue of granule biogenesis (blue vs. green; Fig. 1D) but not acidification by CHGBΔMIF excluded the hypothetical CHGB-less anion conductances because granular biogenesis and delivery was restored. Our data thus pinned down the CHGB subunit *per se* to the ion shunt conductance. Another question is whether such a CHGB+ conductance could conduct cations. Were it cationic like what was proposed for lysosome acidification,[68] it would probably be a K^+^ channel (BK or Kir) as suggested before.[41, 45] However, as far as we know, none of the known K^+^ channels need CHGB to function [69]. Other channels like Na^+^ or Ca^2+^ do not show such a requirement, either [70, 71]. Additionally, our whole-cell recordings of anionic currents through CHGB+ channels showed that even if any of the assigned cation channels were present in DCSGs, their contributions to ion flux were much smaller (<5%) than that of the anion channels. These considerations strengthen CHGB being the kernel of the anion shunt pathway. The CHGB+ channels may be made of CHGB alone or of CHGB plus other subunits. Our data cannot distinguish the two in SGs, yet.

### Dominant negativity of CHGBΔMIF excludes transient CHGB+ channels and suggests hetero-oligomerization with wild-type CHGB

A third type of CHGB+ conductances we suspected to exist represents those that were activated by wild-type CHGB (transiently CHGB+) and then stayed active after CHGB fell off (CHGB-*less*). This transient CHGB+ channel could be excluded when we investigated the dominant negative effects of CHGBΔMIF by assessing the impact of its overexpression in the presence of WT CHGB, instead under the influence of knockdown effects (Fig. 2A). A neuropeptide Y (NPY)-ClopHensor fusion protein, which by default is delivered to SGs, was co-expressed to substitute DND-160.[67] An *in situ* standard curve was prepared with the NPY-ClopHensor in SGs (Fig. 2B). Comparison of mean granular pH shows that CHGBΔMIF overexpression caused strong granule deacidification (pH 6.8) whereas WT CHGB did not (pH∼5.4). A change of 1.4 pH units was stronger than that from the DND-160-based measurements (Fig. 1E), probably because DND-160 was not retained well in SGs of near-neutral or alkaline pH and gave out weaker signals. Because WT CHGB was present in all CHGBΔMIF-expressing cells, the hypothesized transient CHGB+ channels, if indeed existent, should be activated, remain active and maintain granule acidification.

**Figure 2.**
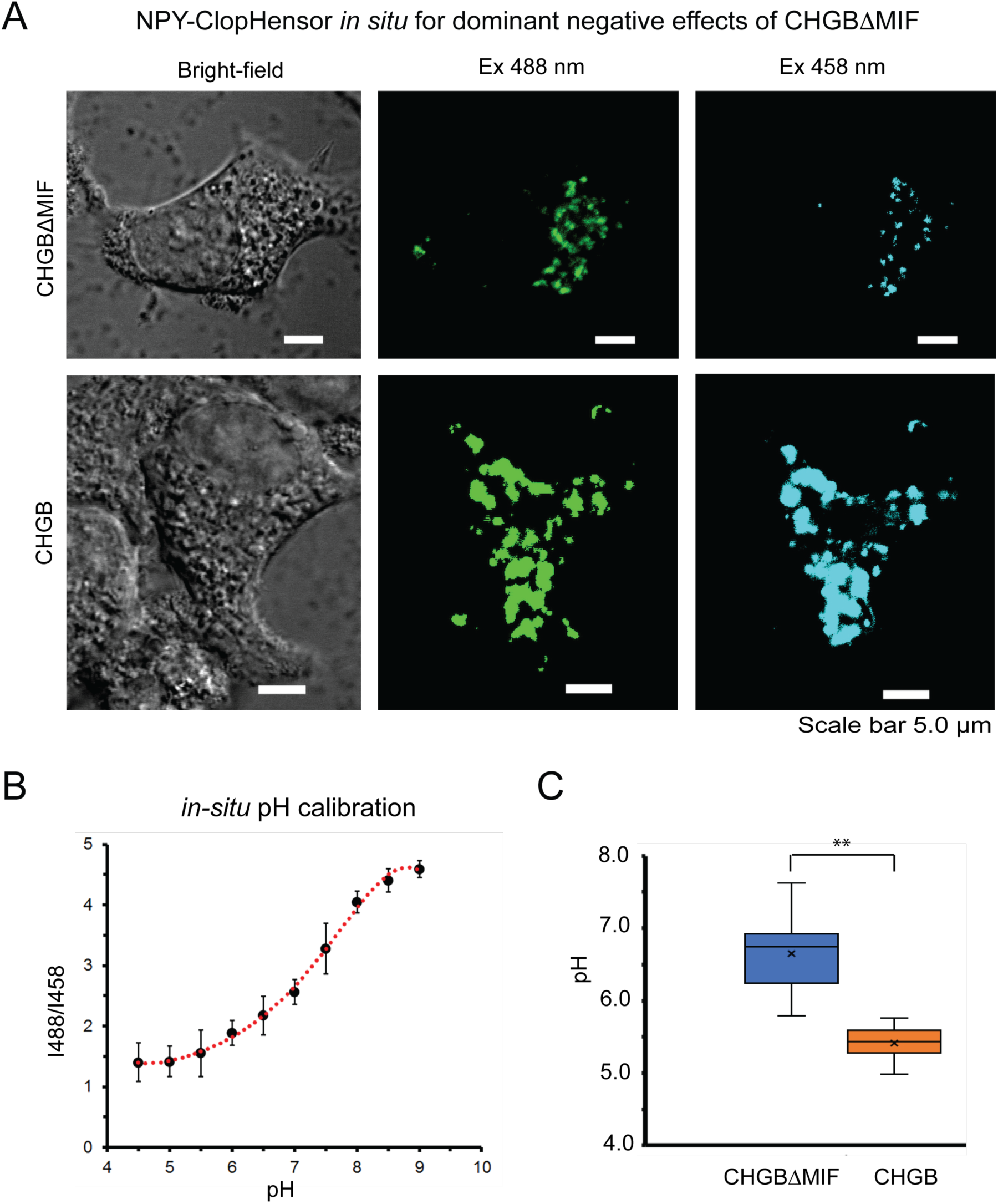
Dominant negative effects of CHGBΔMIF in granule deacidification. (**A**). NPY-ClopHensor was co-expressed with CHGB or CHGBΔMIF in INS-1 cells, which were imaged in a Zeiss LSM710. Typical bright-field DIC images (left column) and fluorescence images at 510 nm with dual excitation at 488 nm (middle column) and 458 nm (right column) were presented. (**B**). A typical *in situ* pH calibration curve in cells expressing NPY-ClopHensor. Experiments were repeated four times. Non-linear regression (polynomial function) was used to fit the data. The obtained function was used to determine the pH levels from the I_488_ / I_458_ ratios measured from individual granules randomly picked out of paired images of each cell. Dozens of cells were randomly selected. Error bars: *s.d*., n=4. (**C**). Average intragranular pH values in cells co-overexpressing NPY-ClopHensor with CHGB (brown; pH ∼5.4) or CHGBΔMIF (blue; pH ∼6.7). **: *p* < 0.01 Errors: *s.d.*, n=3.

The CHGB+ channels made of CHGB only or CHGB plus other partners may both contribute to the anion shunt pathway. They are not separated from each other in SGs because many different proteins are packed in each SG. Both should be inactivated by CHGBΔMIF, probably through oligomerization (Figs. 2A & 2C) because stable CHGB functional units are dimeric and a functional channel was predicted to be a tetramer.[51, 52] A cryo-EM structure of bovine CHGB (bCHGB) dimer revealed an extensive interface for subunit-subunit interactions and enough ectodomains for CHGBΔMIF and WT CHGB to interact. Competition of CHGBΔMIF for the CHGB-binding site must be strong and complete to cripple all resultant hybrid channels. Given that both native and recombinant WT CHGB alone can reconstitute functional Cl^-^ channels, it is reasonable to expect that even in the presence of other subunits, the CHGB-CHGB interactions remain to be the core of highly selective Cl^-^-conducting pores. The oligomerization hence provides a reliable mechanism for strong dominant negative effects.

### CHGB loss impaired proinsulin-insulin conversion in INS-1 cells

Given the tight coupling between cargo maturation and granule acidification, we next inquired about the effects of CHGB loss on the cargo molecules. As an example of small protein cargos, we analyzed semiquantitatively proinsulin-to-insulin conversion in INS-1 cells treated under four different conditions as outlined in Fig. 1B. As a negative control, we examined a key vATPase subunit, ATP6V0A2, by western blotting (WB), and found that its expression level was the same in all four conditions (Fig. 3A), strengthening our earlier conclusion that CHGB loss-induced deacidification is not an indirect result of reduced vATPase activity. It also agreed with the high specificity of the CHGB siRNAs and the fact that CHGB is not required for VATPase activity elsewhere. [51]

**Figure 3.**
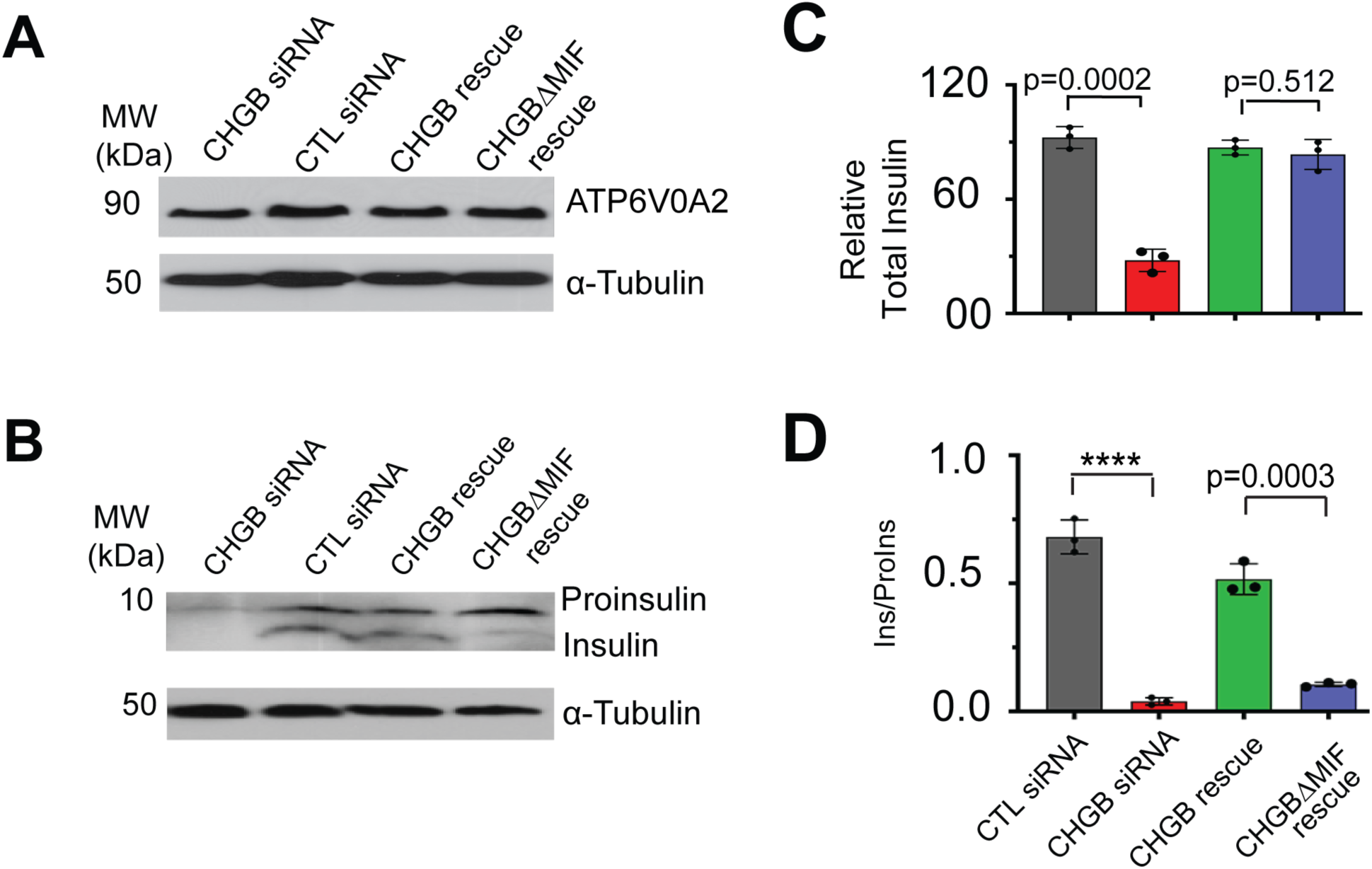
CHGB for insulin maturation in INS-1 cells. (**A**). Western blots of ATP6V0A2 (vATPase subunit) from cells treated in four different conditions as in Fig. 1B. (**B**). Western blots of mature insulin and proinsulin from lysates of cells in four conditions. Loading controls of α-tubulin were used for band quantification of those in **A & B**. (**C&D**). Normalized total insulin (mature insulin + proinsulin; **C**) and insulin maturation (mature insulin / proinsulin ratio, labeled as Ins / ProIns; **D**) in four different conditions. In **C**, total insulin in each was normalized against the total amount from cells treated with CTL siRNAs (black). Error bars: *s.d*. n=3. **: *p* < 0.05; ***: *p* < 0.001; ****: *p* < 0.0001; *ns*: not significant.

Insulin maturation should be impaired (or delayed severely) when a higher granular pH decreases activities of prohormone convertases 2 and 3 (PC2/3) and in turn reduces production of mature insulin. Western blotting of proinsulin and mature insulin (Fig. 3B) showed that CHGB knockdown reduced total insulin (proinsulin + mature insulin) due to reduced granule biogenesis, which was restored by overexpressing CHGB or CHGBΔMIF (Figs. 3B-C). The total insulin is thus determined by cell’s capacity in granule biogenesis (green and blue in Fig. 3C)[30, 66], and was reduced by ∼65% in CHGB knockdown cells (red in Fig. 3C), to the same extent as the reduction in granule numbers (Supplementary Fig. 1B). Notably, CHGB knockdown caused much more severe loss of mature insulin (> 90%, Figs. 3B & 3D) and drove the insulin / proinsulin ratio (Ins/ProIns) close to nil (red in Fig. 3D), indicating more than 90% PC2/3 activity in the leftover SGs were suppressed when luminal pH arose by ∼0.7 units (red Fig. 1G). As expected, overexpressing CHGB restored insulin maturation (>80% of CTL), but CHGBΔMIF failed the rescue (∼10%; green vs. blue in Figs. 3D & 1G). CHGBΔMIF over-expression rescued granule biogenesis and thus the delivery of PC2/3 and proinsulin into SGs (green bar in Fig. 3C), but it failed to restore granule acidification and the production of mature insulin when PC2/3 were suppressed substantially by the pH change.[22, 25]

### CHGB+ channels for filling monoamine-secretory granules in PC-12 cells

Next, we examined if the same mechanism holds for a second type of SGs that carry small organic compounds. We probed CHGB knockdown in PC-12 cells, which contain similar protein machineries as the chromaffin cells and utilize the H^+^ gradient to load dopamine (DA) into SGs (Fig. 4A). As before, WB of ATP6V0A2 found no change (Fig. 4B). We then measured mean granular pH from cells treated with the same four protocols (Fig. 1B). Typical cell images are showed in Supplementary Fig. 4. CHGB loss decreased the average number of SGs by ∼50% (Supplementary Fig. 1C and row 2 in Supplementary Fig. 4), which was reversed by overexpressing CHGB or CHGBΔMIF (rows 3 & 4 in Supplementary Fig. 4). CHGB loss raised the mean granular pH by ∼0.9 pH units (Figs. 4C & 4E), which was restored to ∼5.5 by wild-type CHGB, but not by the nonconducting mutant, CHGBΔMIF (pH ∼6.5; Figs. 4D& 4E). These results phenocopied the treated INS-1 cells, suggesting that the two different types of SGs employ the same CHGB+ channels for anion shunting and granule acidification.

**Figure 4.**
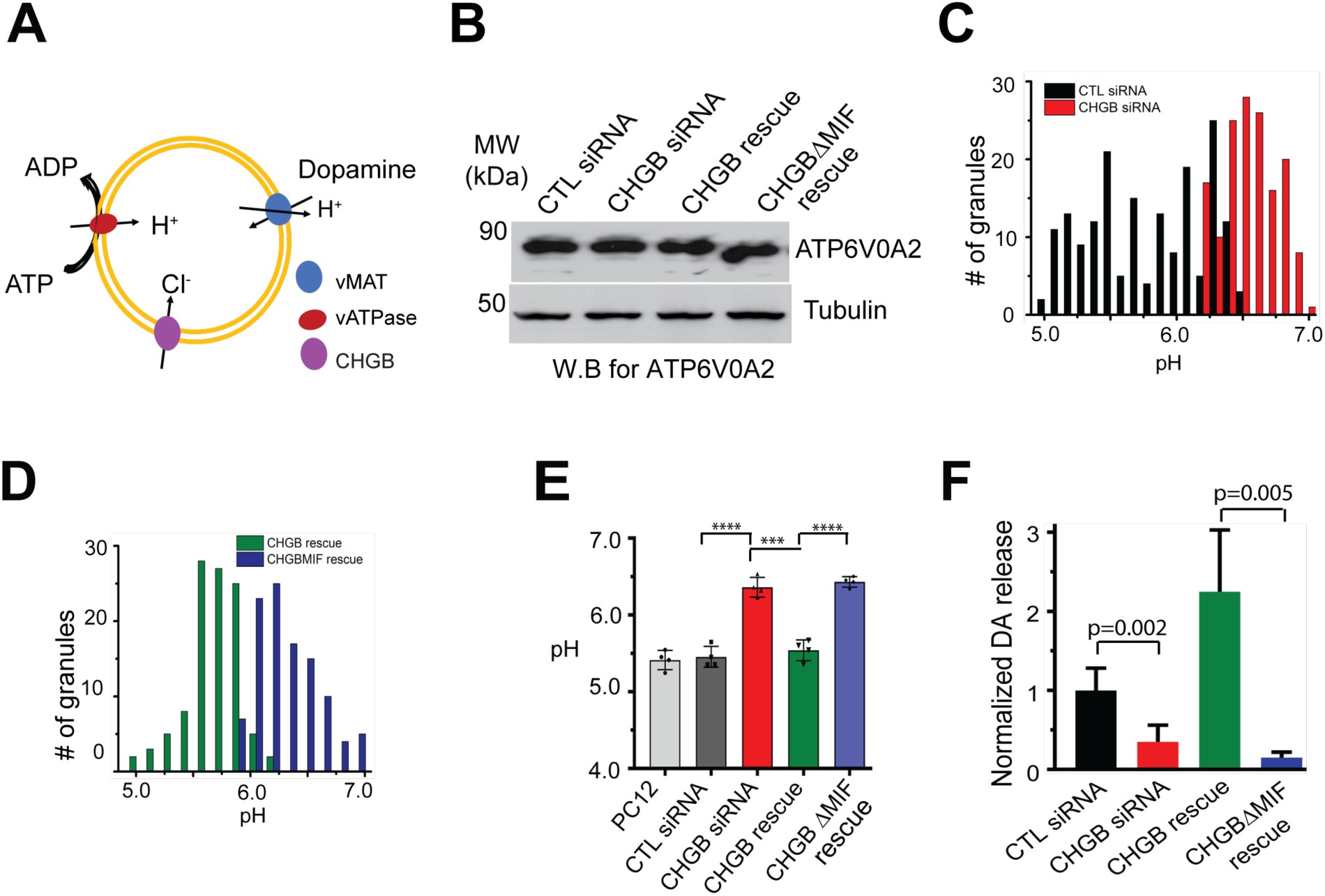
CHGB for normal granule acidification and dopamine loading in PC-12 cells. **(A**) Schematic drawing for H^+^ pumping and Cl^-^ influx that drive granular acidification and H^+^-coupled dopamine (or epinephrine / norepinephrine) loading into granules by vMAT1. (**B**). Western blot of ATP6V0A2 in four differentially treated cells using tubulin as a loading control. (**C**). Distribution of granule pH values from cells transfected with control (black) or CHGB siRNAs (red). (**D**). Histograms of granule pH readouts from CHGB knockdown cells rescued by overexpression of CHGB (green) and CHGBΔMIF (blue), respectively. DND-160 was used. Typical images are showed in Supplementary Fig. 4. (**E**). Average granular pH of cells treated in four different conditions and control PC-12 cells from three different repeats. Errors: *s.d*., n=3. (**F**). Relative dopamine quantities released from the same numbers of depolarization-activated cells treated in four different conditions and measured with an ELISA kit. Standard *t*-test for **E** and two-tailed Welch’s *t*-test for **F**. **: *p* < 0.05; ***: *p* < 0.001 (Errors: *s.d*., n=3).

We then evaluated the effects of granule deacidification on dopamine (DA)-loading in SGs. A vesicular monoamine transporter (vMAT1) pumps cytosolic DA into SGs by dissipating the proton gradient (Fig. 4A).[72] Deacidification by ∼0.9 pH units weakened the driving force for vMAT1 by ∼8 folds and should decrease the DA content in readily releasable granules (RRGs) significantly, ∼90% if no compensation occurred. Using an ELISA kit to measure normalized DA amounts released by RRGs upon stimulated granule exocytosis, we observed that CHGB knockdown decreased RRG DA content by ∼80%, which was recovered by WT CHGB (∼200%, Fig. 4F). The overshooting in the CHGB rescued cells probably stemmed from enhanced granule biogenesis (Supplementary Fig. 3). In contrast, CHGBΔMIF did not restore the DA release (Fig. 4F), but instead drove the RRG DA content to a level (5-10 % of control) even lower than that (∼25%) of the knockdown cells, showcasing again the dominant negative effects. Consistently, DA content measured by carbon fiber electrodes was decreased substantially (by ∼60%) in chromaffin cells from the *Chgb^-/-^* mice.[60-62, 73]

### Native CHGB+ channels support granule acidification in primary pancreatic β-cells

We next tested the conservation of the above molecular mechanism in primary neuroendocrine cells in a mouse model. Prior analyses showed that *Chgb-/-* mice exhibited reduced catecholamine content in chromaffin cells, decreased insulin secretion from isolated islets, abrogated 2^nd^-phase insulin release after glucose challenge, and uplifted serum proinsulin (hyperproinsulinemia).[60-62] At that time of publication, these four categories of phenotypes were apparently unrelated to each other and quite puzzling in view of CHGB’s Ca^2+^-binding capacity and its contribution to proteolytic peptides. However, the CHGB+ channel shed a new light.[61] To evaluate its effects, a strain of *Chgb*^-/-^ mice from EMMA (#10088, URL: https://www.infrafrontier.eu/) were raised in the Animal Care Service facility at the University of Florida (UF) School of Medicine. An animal protocol (#201709886) was approved at UF and was followed verbatim. Genotyping was performed by PCR of tail nips (Supplementary Fig. 5). Pancreatic islets were acutely isolated from 8 - 10 weeks old mice. Age-matched male and female littermates were processed separately.[74] Fig. 5A shows representative images of isolated islets (left). WB confirmed no CHGB protein in *Chgb-/-* mice (right). Intragranular pH was then measured in either isolated islets (Figs. 5B-D) or β-cell monolayers grown out of fresh islets (Figs. 5E-F).

**Figure 5.**
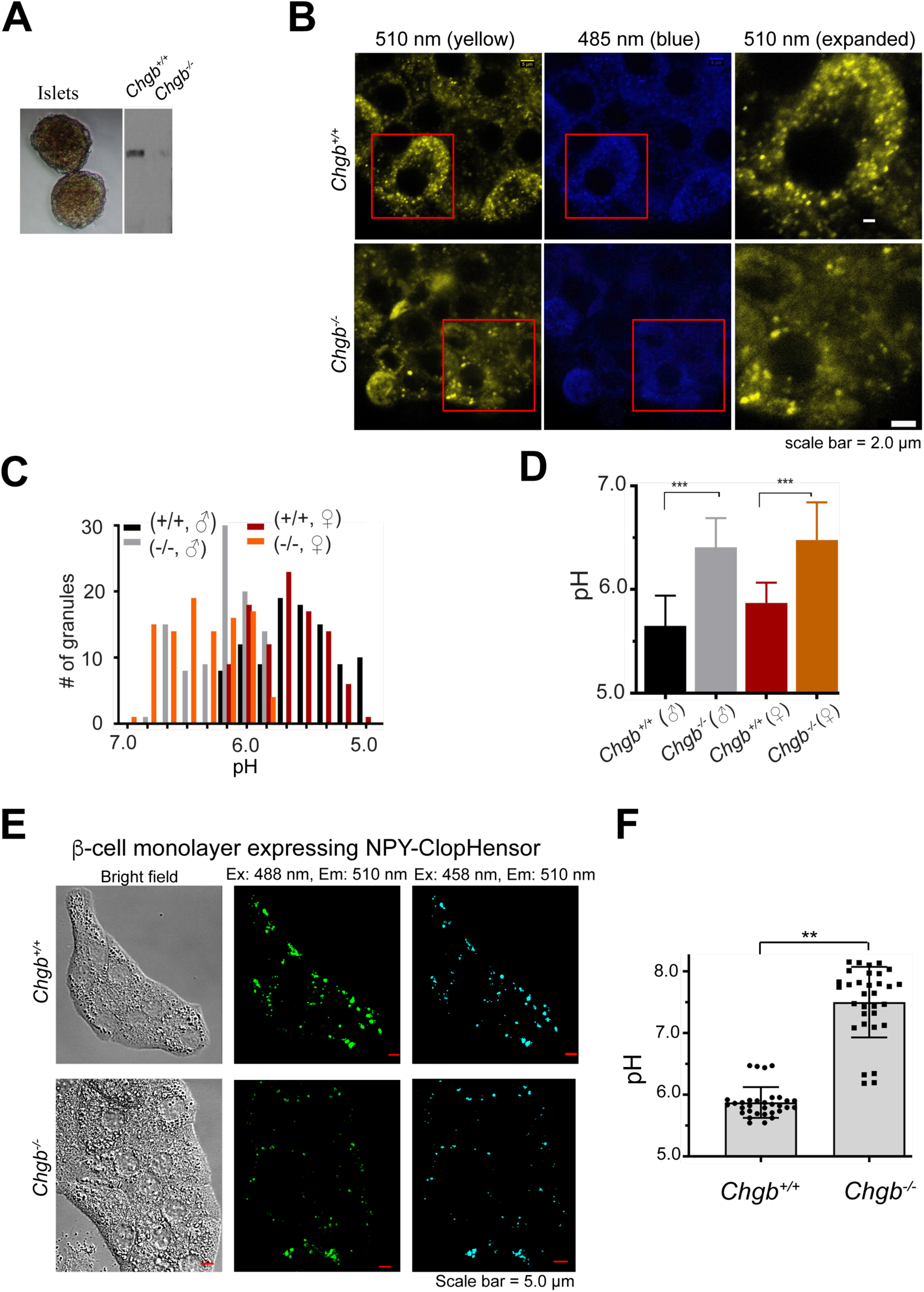
CHGB+ channels function for normal granule acidification in primary β-cells. (**A**). Two typical pancreatic islets from knockout mice (left) and western blots of CHGB from liver tissues of a wild-type (*Chgb^+/+^*) and a CHGB-knockout (*Chgb^-/-^*) mouse. (**B**). Typical ratiometric images of pancreatic β-cells in individual islets from *Chgb*^+/+^ and *Chgb^-/-^* mice. Typical images out of 10 repeats were shown. Excitation: 410 nm; emission: 485 nm and 510 nm. A small area (red window in the left) in a 510 nm image was expanded to show granules in the rightmost column. β-cells from *Chgb^-/-^* mice have fewer bright granules (above background) and weaker signals at 510 nm due to deacidification. (**C**). Histograms of intragranular pH values measured from granules in islet cells from both male and female mice of different genotypes. All mice were ∼9 weeks old. (**D**). Average pH values for four different groups of mice as in **C**. **: *p* < 0.05; ***: *p* < 0.001 (Errors: *s.d.*; n=4). (**E**). Typical ratiometric images in monolayers of β-cells grown out of islets from *Chgb*^+/+^ and *Chgb^-/-^* mice. After ∼7 days, monolayers were transfected to express NPY-ClopHensor and imaged after 48-72 hours. Excitation: 488 nm (green) and 458 nm (cyan); emission: 510 nm. β-cells from *Chgb*^-/-^ mice have a lower number of granules than the wild-type cells. (**F**). Average intragranular pH measured from ClopHensor in monolayer β-cells from *Chgb*^+/+^ and *Chgb*^-/-^ mice. **: *p* < 0.05. Data were pooled from three repeats. Error bars: *s.d.*, n=35.

Compared to WT cells (top row in Fig. 5B), *Chgb*^-/-^ β-cells (first 2 - 3 layers of cells underneath the islet surface) contained fewer well-stained granules (bottom, Fig. 5B), suggesting reduced biogenesis (Supplementary Figs. 1B-C; Figs. 1F, 3C). Histograms of intragranular pH values showed a significant shift to higher numbers in *Chgb*^-/-^ β-cells from both male and female mice (Fig. 5C). The average intragranular pH in *Chgb*^-/-^ β-cells was ∼0.7 pH units higher than that of the WT cells (∼5.6 to ∼6.3 for male, and ∼5.7 to ∼6.4 for female; Fig. 5D), the same as the data from CHGB knockdown cells (Fig. 1G). As predicted, loss of CHGB+ channels *in vivo* results in persistent granule deacidification in native neuroendocrine cells (Figs. 5C-D). Deacidification expectedly delays proinsulin-to-insulin conversion, increases proinsulin content in isolated *Chgb*^-/-^ islets, and results in hyperproinsulinemia but a similar basal level of serum insulin due probably to compensation.[61]

DND-160 staining of enzyme-liberated islets suffered from potential complications from uneven staining of multiple layers of cells and higher background noises from stained connective tissues. To resolve this concern, we grew β-cell monolayers from acutely isolated islets in substrate-coated culture dishes [75] and then transiently expressed NPY-ClopHensor for granular pH measurements (Fig. 5E). Compared to data from wildtype cells, the average granular pH increased by ∼1.6 pH units (5.8 to 7.4) in *Chgb*^-/-^ cells, greater than the change of ∼0.7 pH units in DND-160-stained islets (Fig. 5F vs. 5D), probably due to higher sensitivity of the ClopHensor in the alkaline pH range (pH>7.0, Fig. 2B) and lower DND-160 signals in SGs of neutral or alkaline pH. Nonetheless, these results corroborated very well the persistent granule deacidification in cells inside isolated islets (Fig. 5D).

### CLC-3/5 or CACCs appear nonessential to the anion shunt conductance

With the CHGB+ channels being functional in SGs, we could evaluate possible contributions of CLC3 or other proteins to anion shunting. In the *Chgb*^-/-^ mouse tissues, WB data suggested that pancreatic CLC-3 expression was not decreased but was instead increased by ∼2 folds (Fig. 6A). The persistent granule deacidification in *Chgb^-/-^* β-cells therefore ought not to have been derived from lower CLC-3 expression, or loss of CLC-3 functions. Further, we compared the effects of knocking down CLC-3, CLC-5 or the TMEM16 family calcium-activated chloride channels (CACCs), ANO-1/2 (TMEM16A/B) to be specific, in cultured INS-1 cells by using specific siRNAs. None of the four knockdowns resulted in significant granule deacidification as CHGB siRNAs did (Fig. 6B). WB was performed to confirm that these siRNAs were effective and specific for their targets and did not suppress CHGB expression. Similarly, CHGB siRNAs did not change CLC-3 or CLC-5 expression, either.[51] All these results convergently support the conclusion that the granule deacidification in CHGB knockdown cells was not an indirect result from loss of CLC-3/5 or CACCs, but was a direct one from loss of CHGB in SGs. This conclusion accords well with our findings that only the CHGB+ channels have the capacity of the anion shunt conductance and the few cation channels (CHGB-less) assigned to the SGs could not replace CHGB.

**Figure 6.**
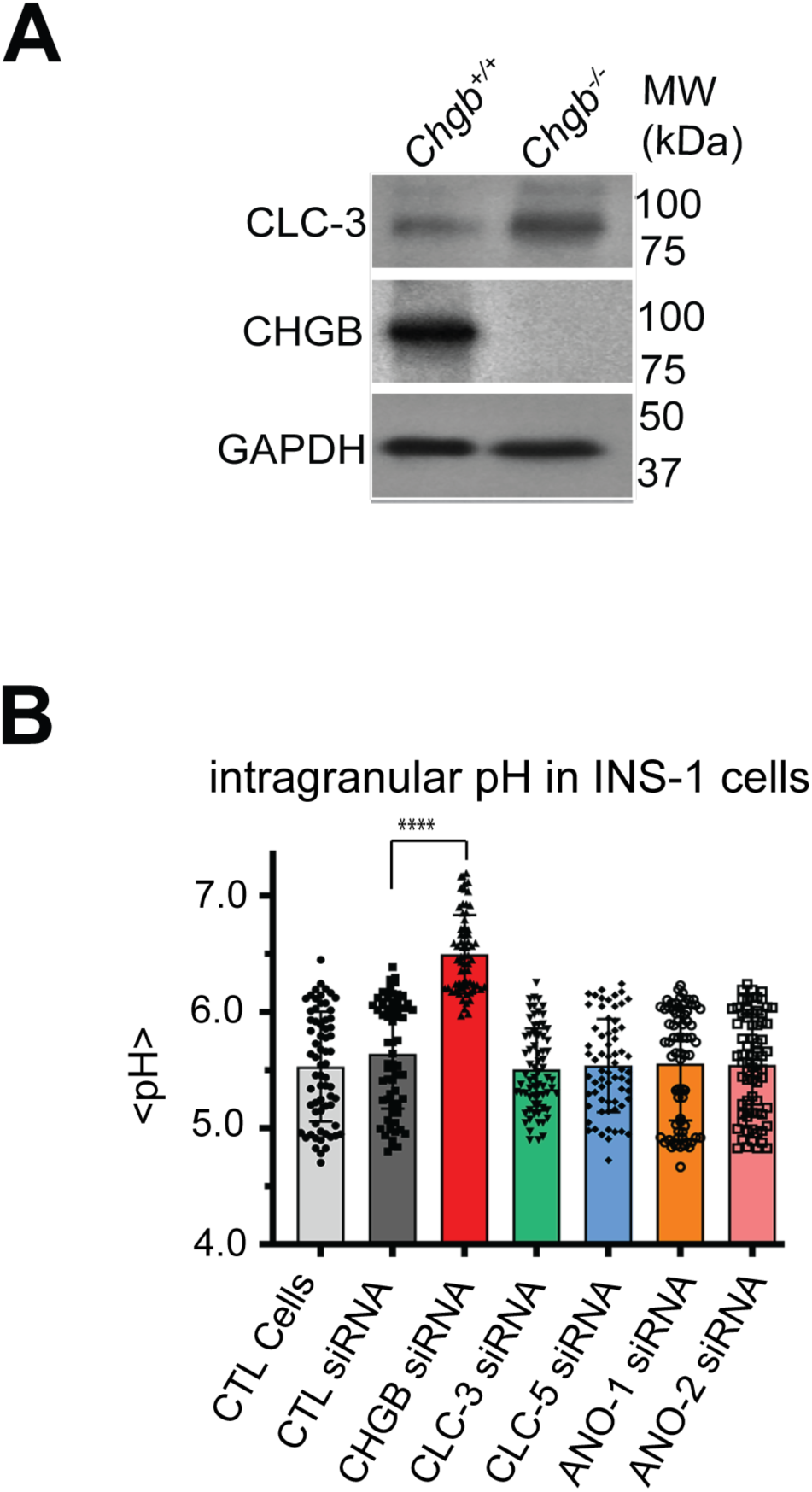
CHGB+ channel functions are independent of CLC-3/5 or ANO-1/2. (**A**). Western blotting of CLC-3 and CHGB in pancreatic tissues from *Chgb*^+/+^ and *Chgb*^-/-^ mice. GAPDH as loading controls. (**B**). Knockdown of CLC-3, CLC-5, TMEM16A (ANO-1) or TMEM16B (ANO-2) did not change granule acidification in INS-1 cells. Average pH values were calculated from hundreds of stained granules pooled from three repeated experiments. **: *p* < 0.05, n ∼150. Three repeated experiments yielded the same results and were pooled together for statistical analysis.

## DISCUSSION

Our observed effects from dismantling the CHGB+ channels match exactly what the long-sought anion shunt conductances do in two typical types of SGs in cultured and primary neuroendocrine cells, providing a well-grounded solution to a nearly 50-year-old question in eukaryote Cell Biology (Fig. 7A). As the kernel of the CHGB+ channels, CHGB loss causes granule deacidification in INS-1, PC-12 and *Chgb-/-* beta-cells (Figs. 1G, 4E, 5D & 5F), slowdown of proinsulin-insulin conversion (Fig. 3), and diminution in dopamine loading (Fig. 4F). The CHGB+ channel, although being dominated by an unconventional dimorphic protein, represents a new biochemical signature of the SG membranes. The dominant negative effects of CHGBΔMIF stem from its strong interaction with WT CHGB, which diminishes functional CHGB+ channels (Fig. 2). Loss of the CHGB+ channels disables anion shunt currents and drives deacidification and retards cargo maturation severely in both insulin- and monoamine-SGs (Figs. 1A, 7B & 7C).

**Figure 7.**
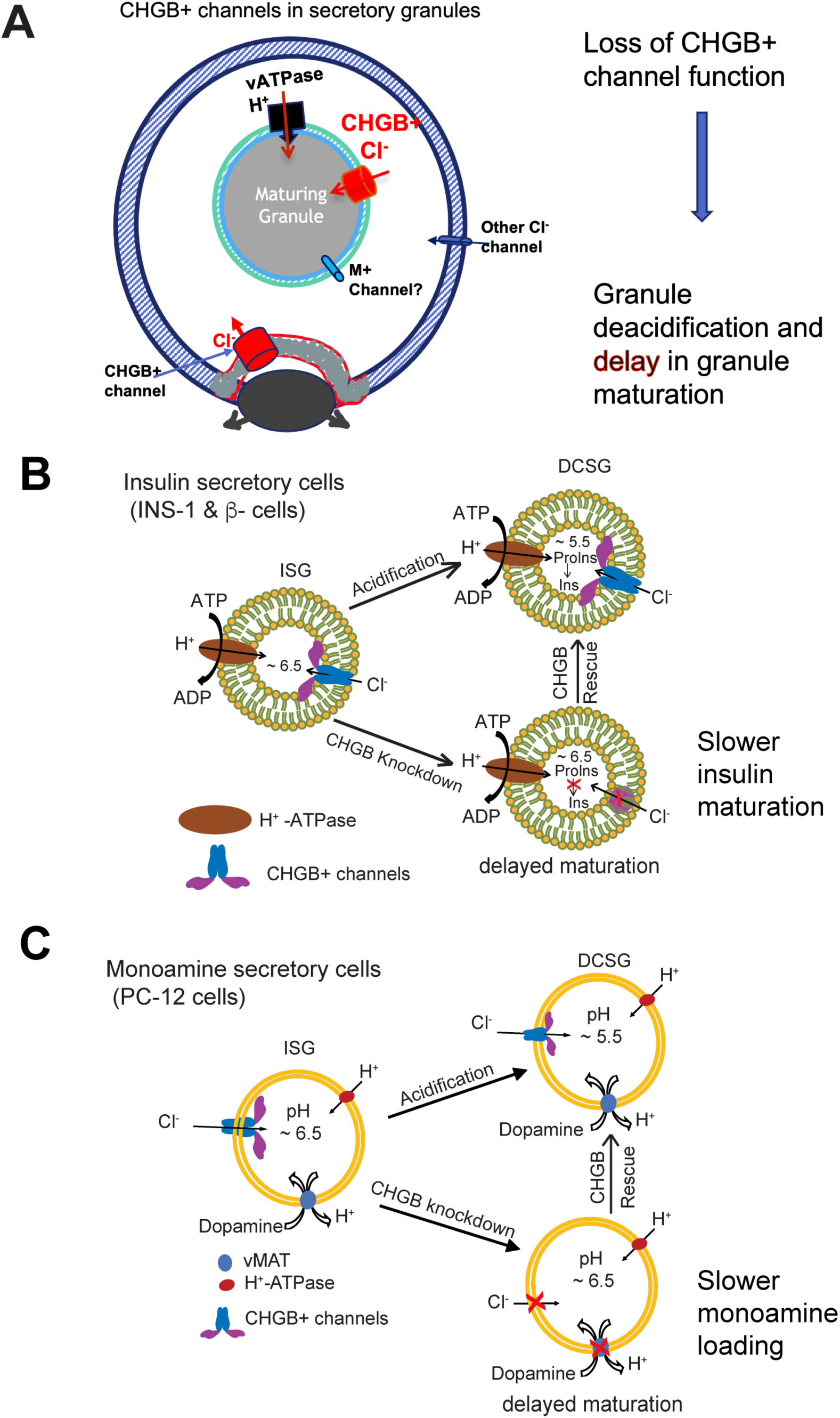
Molecular mechanisms for CHGB+ channels in regulated secretion. (**A**). CHGB+ channels in the granules are functional for the anion shunting pathway and facilitate continued proton pumping by vATPase. Its loss will slow down H^+^-pumping and delay cargo maturation. (**B**). CHGB+ channels for normal acidification of insulin-secretory granules. CHGB knockdown leads to the loss of CHGB+ channel function, impairs normal acidification and slows down proinsulin-to-insulin conversion. CHGB overexpression restores acidification and insulin maturation, but not the deletion mutant CHGBΔMIF. (**C**). In monoamine-secretory cells (PC-12 or chromaffin cells), loss of CHGB+ channels impairs granule acidification and diminishes driving force for H^+^-coupled transport of catecholamines (*e.g.,* dopamine) into secretory granules. Overexpression of CHGB, but not CHGBΔMIF, rescues the defects.

Our data do not support a major contribution to SG ion balance of either cation shunt channels or CHGB-less Cl^-^ channels / transporters, of which CLC-3/-5 and ANO-1/2 represent only a small fraction (Fig. 6), probably because CHGB-less anion transporters or channels are either absent in the SGs (like ANO-1/2) or present in a quantity (CLC-3 as a possible example) too low to be as effective as the large-conductance CHGB+ channels.[37] Because of the ample ion flux needed for effective acidification, an electrogenic transporter in a low copy number cannot satisfy the physiological need for charge neutralization in a short time. For example, the proposed CLC-3 splicing variants might fall into this category.[49] More broadly, our results exclude three groups of hypothesized CHGB-less anion channels: (i) Channels whose transcription and/or expression, but not functions, requires CHGB (as a signal); (ii) Channels whose delivery into SGs, but not function, relies on CHGB (as a chaperone), and (iii) Channels that are transiently activated by CHGB (as an activator), but stay active after CHGB falls off. We thus expect that the CHGB+ channels remain functional when migrating from ER/Golgi membranes through the regulated secretory pathway to appear on the cell surface.

Regarding the proposed channels made of CHGB and other subunits, we have very little information on what roles the non-CHGB subunits play and whether they differ when CHGB is in ER, trafficking vesicles, Golgi, TGN, SGs, RRGs, or the plasma membranes. From the available data, we expect that these subunits support, instead of blocking, anion shunt currents in SGs. Analyzing the modulation of native CHGB+ channels in SGs or on the cell surface after their release may provide more insights on whether the non-CHGB subunits really modulate CHGB+ channels and whether they make indispensable contributions to other cellular processes, such as CHGB distribution between soluble and membrane-integral states, CHGB recycling through endosomal compartments, etc. Deciphering their genetic identity would bring a strong boost for further investigations.

Our data lend little support for CLC-3 to be a key part of the anion shunt pathway. Given that, how can the roles of the CHGB+ channels be reconciled with the phenotypes of the *Clcn3-null* mice?[33-35, 49] One potential scenario comes from the possible roles of CLC-3 in the reuptake of CHGB+ channels from the cell surface and then the channel redistribution.[76] If so, without CLC-3 and over a longer period the CHGB+ channels in primary beta-cells might not return to SGs well or might be modified to become nonfunctional as a cellular adaptation to CLC-3 loss, both of which would deter granule deacidification to certain extents. Another possible scenario hinges on CHGB’s dimorphic nature --- the protein existing in two forms. If CLC-3, which we assume is present in SGs in small quantities,[49] facilitates CHGB insertion into membranes, its loss over a long time of cell line or mouse development could keep more CHGB preferentially in its soluble state. Such a shifted distribution would render a major fraction of SGs to lose anion shunting and delay acidification, sufficient to cause ∼44% reduction of H+-pumping due to positive potentials high enough (∼80 mV) to block H^+^ flux and ∼80% loss of granule exocytosis (lost fusion events)[34] due to a major reduction of mature RRGs and partial suppression of Ca^2+^ release from SGs or influx from outside right before granule release. Other possibilities might be considered when more experiments will be performed to test the above two in *Clcn3-/-* cells or mouse models in the future.

Dimorphism is an intrinsic property of CHGB, which accords well with the prior data obtained with soluble CHGB fractions.[59] From ER to Golgi, a fraction of nascent CHGB will likely be partitioned into membranes and form anion channels, which should make these membranes highly permeable to Cl^-^. After SG release, hundreds of CHGB+ channels delivered to the surface per cell make the plasma membrane highly Cl^-^-permeable. [51] The CHGB in the plasma membrane will be endocytosed in a couple of hours and resorted back into TGN, nascent SGs, or lysosomes, and probably make these organellar membranes permeant to Cl^-^ as well.[50] Given the importance of CHGB+ channels in granule maturation, it makes sense that CHGB is genetically associated with type 2 diabetes (Supplementary Table S1),[77] and NATURE has evolved backup solutions to deal with CHGB loss. The hypothesized compensatory effects may explain the milder phenotypes in the *Chgb* or Chga*/Chgb* double knockout mice. In *Chgb*^-/-^ mice, CHGA was reported to be upregulated, probably a good substitute for CHGB’s function in granule biogenesis [66]. In Chga/Chgb double knockout mice, compensation from other gene products allows granule biogenesis to be seemingly normal in neuroendocrine cells.[73]

The disparate phenotypes exhibited by the *Chgb*-null mice can be unified around the CHGB+ channel functions.[60-62] In these mice, no other channels were able to restore the disabled anion shunt pathway, leading to persistent deacidification. A time-integral of diminished vATPase activity (say ∼5% of WT) may be enough to slow down cargo maturation over an extended period, which can make *Chgb*^-/-^ mice able to compensate for plasma insulin production by inducing a higher number of SGs (possibly by higher CHGA production), leaving the mice seemingly healthy under well-controlled feeding conditions. The slowdown of proinsulin-insulin conversion leads to high plasma proinsulin and the loss of the 2^nd^ phase insulin secretion upon glucose challenge because this phase happens in a short time (∼20 minutes). From this same view point, a relatively normal level of phase 1 insulin release is attained when the SGs have a longer time to become maturated and the number of SGs rise up in the *Chgb-/-* β-cells, but a significantly lower level of phase 2 insulin release from nascent granules that failed to mature properly in a limited duration (20-30 minutes).[61] Understanding these mechanisms, especially the fast and slow aging of ISGs and DCSGs and the release or recycle of aged DSCGs, awaits future studies.

CHGB channels may embody significant clinical implications. The main phenotypes of *Chgb*^-/-^ mice suggest a bonding tie of the CHGB+ channels to missense variants genetically associated with type 2 diabetes in humans, [60-62] as listed in Supplementary Table S1[77], and to mutations strongly linked to cardiac diseases, neuroendocrine cancer or neurodegenerative diseases [77-81]. For example, overexpression of CHGB and CHGA together with HSP90 was observed in almost all metastatic human neuroendocrine cancer tissues showing metabolic defects,[79] but their functional significance remains mysterious. If a causative relation between elevated activities of CHGB+ channels and continued growth or survival of neuroendocrine cancer cells can be established, these channels will become a promising target for innovative molecular therapeutics.

Our studies may be expanded in three different aspects. One is to measure directly ion flows across granule membranes inside cells. With the SGs too small for direct patch clamp recordings, a highly specific sensor for Cl^-^ or transmembrane potential could be employed to monitor ion flows during granule maturation. But targeted delivery of highly specific dyes for Cl^-^ has been difficult to achieve. The second is to obtain highly purified SGs and fuse them for direct patch recordings in vesicles or planar lipid bilayers. High biochemical purity of cellular membrane preparations and fusion of granular membranes are not easy to control. The third is to identify possible partners of CHGB in the anion shunt pathway, which will require a new set of experiments. With new technical developments, these three aspects will become feasible and may enrich our understandings significantly.

In a broader scope, functions of the CHGB+ channels are expected to be universal in all CHGB-expressing organisms, which include all vertebrates.[13] Lower-order eukaryotic organisms, especially unicellular ones, lack CHGB orthologs and are expected to utilize substitute molecules to establish an ion shunt pathway with vATPase in SGs. The genetic identity and biochemical properties of these substitute conductances are yet to define.

## CONCLUSION

The CHGB+ channels in the secretory granules of regulated secretion serve the functions of the anion shunt pathway that is coupled tightly to granule acidification and cargo maturation. Such an *in-situ* function of CHGB explains parsimoniously all our observations as well as the once puzzling phenotypes diagnosed in *Chgb*-null mice. All data together disfavor significant contributions of cation conductances or CHGB-less anion channels to ion shunting because these conductances, even if present in SGs, fail to restore granule acidification and cargo maturation in face of CHGB loss. The well-conserved CHGB orthologs in vertebrates thus constitute, alone or in conjunction with other subunits, a new type of intracellular anion channels that may function in membranous compartments upstream or downstream of regulated secretion in various CHGB+ cells, including neurons and neuroendocrine cells. The dimorphic nature of CHGB accords with the previous observation of soluble CHGB in biochemical separation. The proposed indirect effects of CLC-3 on CHGB sorting and redistribution could explain the observations in *Clcn3*-knockout mice and will need further validation. Potential functions of the CHGB+ channels in ER and Golgi, budding of nascent granules, homotypic fusion of and content resorting in ISGs, DCSG aging, recycling compartments, etc., await future investigations.

## METHODS

### 1. Molecular cloning of CHGB and its mutants in pFastbac1

Preparation of all constructs used the same procedures as described before.[52]

### 2. Knockdown of CHGB in neuroendocrine cells and its rescue by CHGB or CHGBΔMIF

Details for cell culture and transfection with siRNAs as well as rescue by relevant plasmid DNAs were described before. [51]

### 3. Detection of insulin and proinsulin in INS-1 cells

INS-1 cells were transfected with 100 nM CTL siRNAs or CHGB-targeting siRNAs in a 6-well plate, and 48 hours later, were fed with fresh medium. Ninety-six hours after transfection, cells were collected for analysis. For the rescue experiments, cells transfected with CHGB siRNAs were incubated for 48 hours, washed, and then transfected with cDNAs for wild-type CHGB or its deletion mutant (CHGBΔMIF) cloned in the pcDNA3.1 vector. After two more days the cells were collected for analysis.

To measure insulin maturation semiquantitatively, INS-1 cells were lysed by freeze-thaw cycles as described before.[60] A 12% tricine-SDS-PAGE gel was used to separate mature insulin and proinsulin. The internal loading control was used to correct for small variations. An anti-insulin antibody (L6B10 from Cell Signaling) was used to detect both mature insulin and proinsulin. The band intensity for both mature insulin and proinsulin in the western-blotting images was measured in ImageJ, normalized against the loading control and compared for total insulin that is mature insulin plus proinsulin, and the maturation of insulin was expressed as ratios of mature insulin: proinsulin (Ins / ProIns).[82]

### 4. Western blotting of vacuolar ATPase subunit A2 (ATP6V0A2)

Details were described by Yadav *et al*.[51]

### 5. ELISA for dopamine release from PC-12 cells

On day 0, 3 x10^5^ PC-12 cells were seeded in a 6-well plate. Transfection with CHGB siRNAs was performed on day 1. On day 3, cells were washed and transfected with plasmids carrying cDNAs for wild-type CHGB or CHGBΔMIF. On day 5, cells were collected, and the same number of non-transfected cells were prepared as control. The knockdown cells without rescue were fed with fresh medium on day 3. Dopamine secretion was analyzed as previously described with slight modifications.[83] The cells were washed twice in a low K^+^ buffer (20 mM HEPES-NaOH, pH 7.4, 140 mM NaCl, 2.5 mM CaCl_2_, 1.2 mM MgCl_2_, 5.0 mM glucose, and 4.8 mM KCl). Equal numbers of cells under differential treatments were processed in parallel with a low K^+^ (4.8 mM KCl) or high K^+^ buffer (40 or 55 mM KCl to replace equivalent NaCl in the normal buffer) for 11 min at 37 °C. Afterwards, the media were collected and centrifuged at 10,000 × g for 20 s at 4.0 °C to remove all cells or cell debris. The supernatant dopamine was measured with an ELISA kit (Eagle Biosciences). We performed these experiments four times and observed no obvious toxicity of siRNA or cDNA transfection to PC-12 cells. The measured amounts of dopamine were normalized against that from CTL siRNAs-treated cells. The amounts of total proteins in cell lysates were measured to confirm that the measured dopamine was released from approximately the same number of cells in four different conditions.

### 6. Ratiometric pH measurements in DND-160-stained cells

Although well-established protocols were available, we first tested whether the lysosensor DND-160 was suitable for our measurements. INS-1 cells were transfected to express syncollin-pHluorin fusion protein [64] (a cDNA construct kindly provided by Dr. Herbert Y. Gaisano in the Department of Medicine at University of Toronto, Canada). Syncollin is by default delivered to secretory granules. The pHluorin protein fluoresces well in acidic pH and serves as a good tracer. Forty-eight hours after transfection, the cells were washed with a pre-warmed physiological medium (10 mM HEPES, 120 mM NaCl, 4.8 mM KCl, 2.5 mM CaCl_2_, 1.2 mM MgCl_2_, 24 mM NaHCO_3,_ 3.0 mM glucose, pH 7.4) and incubated with 0.50 µM DND-160 for 5 minutes at 37°C. Afterwards, the cells were rinsed quickly and imaged in a Zeiss LSM 780 confocal fluorescence microscope with a 63 x oil-immersion lens using 458 / 485 nm excitation and 510±10 nm emission for pHluorin and 405 nm excitation and emission at 510 nm and 485 nm for dual emission measurements of DND-160. At the University of Florida and SUNY-Buffalo, Zeiss LSM 800 confocal microscopes were used instead. A calibration curve was obtained for each microscope before ratiometric readings.

INS-1 or PC-12 cells were prepared as described above. Imaging was always done 96 hours after siRNA transfection. Immediately before imaging, cells of similar confluency were washed with the physiological medium and then stained with 0.50 μM DND-160 for 5 minutes with. 0.50 µM bafilomycin added to block vATPase activity. Excess dye was washed away with the physiological medium and paired images of the cells were captured in a confocal microscope. For a double excitation experiment, a 40-x quartz objective lens was used, excitation for DND-160 was set at 340 nm or 380 nm and emission at 510 nm was measured. We utilized a home-assembled Fura-2 imaging system in Dr. Jen Liou’s laboratory at UT Southwestern Medical Center. For every granule in each pair of images, a circular mask of the granule size was used to read average intensity of the granule. In close vicinity to the granule, the same mask was used to read out background signal from four neighboring blank areas. The averaged background signal was subtracted from the averaged granular density, generating the final intensity for the selected granule. The paired images were analyzed side-by-side so that the same areas were selected for each granule. A 340 nm / 380 nm ratio was calculated as the dual excitation readout. For dual emission, the cells were imaged under an LSM 780 (or LSM800) using an apochromat 63x/1.40 Oil DIC objective lens with excitation at 405 nm and emissions at 510 nm (pseudo-colored in yellow) and 485 nm (pseudo-colored in blue). 20 nm apertures were used. After background subtraction, ratios of averaged densities at 510 / 485 nm for individual granules were calculated. The standard curves for dual excitation or dual emission were prepared for each microscope using the DND160 in buffers of different pH values. The same standard curve was used to minimize cell-to-cell variations. Because of the reported difficulty in controlling pH inside the granules for calibrating pH *in situ* [35], we used *in vitro* calibration and measured pH values from hundreds of granules to minimize possible background variations among granules within the same cell or from different cells. Histogram distribution of measured pH values therefore reflected the distributions of the ratiometric measurements based on the calibration curve and may not reflect exactly the *in-situ* pH values. However, the observed trends should reflect well the *in-situ* changes.

To overcome potential limitations in the DND-160-based pH readout, especially in the range of pH variations and the trends of pH changes, positive and negative control experiments, *e.g.,* NH_4_Cl treatment and transfection with control siRNAs, were used to demonstrate the suitable application of the ratiometric measurements from DND-160 as a reliable quantification of the alterations in intragranular pH. Many granules were randomly selected from multiple cells in each condition to overcome small variations from one cell to another and from granules of different sizes or in different states and derive statistically significant and objective comparison. Because of diffraction limit and the possible clustering of multiple granules, it is difficult to determine granule sizes based on the fluorescence spots. To evaluate the possible effects due to variation in the duration of dye-loading, cells stained with DND-160 for 5 minutes were compared with those stained for 30 minutes. No difference in the measured pH-distribution was observed using a longer loading time. It appeared that 5 minutes was sufficient to reach a sufficient level of dye loading and was suitable for our experiments. As a different control, we compared pH measurements using DND-160 staining and ClopHensor expressed *in situ* and found that the two different dyes detected the same trend in granule acidification and deacidification.

### 7. Isolation of pancreatic islets from mice

A UF IACUC-approved protocol (#201709886) was developed by the Jiang lab for experiments with mice. We imported the CHGB knockout mice from InfraFrontier GmbH in Germany. The heterozygous (*Chgb*^+/-^) C57BL/6 strain was marketed as #EM: 10088 in the EMMA deposit with a strain name of CEPD0073_3_D01 and a designation of **C57BL /6NTac-Chgb^tm1(EGFP/cre/ERT2)Wtsi^/WtsiIeg**(www.infrafrontier.eu/search?keyword=CHGB). The genome-wide mutants were first produced by Wellcome Trust Sanger Center and EMMA is a distributor in Germany. The mice were raised from frozen sperms as a heterozygous colony of 3 males and 2 females and shipped to the UF Animal Care Service (UF ACS). After being in quarantine at the UF ACS for three months, the mice were raised inside the animal breeding facility so that environmental conditions were strictly maintained during the whole period of our experiments. Genotyping all offspring was performed by PCR amplification of DNAs extracted from tail nips. Information for the primers used for PCR was obtained from EMMA. Western blotting of full-length CHGB was performed from processed tissues including liver, pancreas, spleen, and kidney, etc. We kept three pairs of heterozygous mice for continuous breeding inside the Animal Breeding Facility. Every time, we used two wildtype (one male and one female) and two homozygous (one male and one female) mice for surgical preparation of pancreatic islets. The mice were dispatched by the Animal Care Service on the day of experiments and sacrificed on the same day. The same type of experiments were always repeated more than 5 times to gain statistical significance and assure reproducibility.

Surgical removal of pancreas from mice followed a standard protocol with some modifications [74, 84]. After being euthanized with CO_2_, a mouse was laid in its supine position and sanitized by spraying with 70% ethanol. Its abdominal cavity was surgically opened to expose small intestine, common bile tubes, pancreas, etc. Cardiac aorta was cut to ensure complete animal euthanization. Under a dissecting microscope, small intestine was repositioned so that the common bile duct followed a line perpendicular to the vertical axis through the head of the mouse. The common bile duct was clamped in a position as close to the small intestine as possible. Two milliliters of Liberase TL (Roche; distributed by Sigma-Aldrich) at 0.9 units / ml were injected slowly using a 27G1/2-gauge needle via a nylon tubing through the bile duct. After the injection, the whole pancreas was gently removed from the animal and stored in a 50 ml conical tube in ice. Usually, all four mice were processed in one session. Each took about 30 minutes. The genetic phenotype and gender of each mouse were marked on the conical tube. After surgery the mouse carcasses were properly disposed of.

After surgery, the conical tubes containing the mouse pancreas were incubated in a water bath at 37 °C for 14 minutes. Twenty-five milliliters of HBSS supplied with Ca^2+^ and Mg^2+^ were added to each tube, and the pancreatic tissues were disintegrated by gently pipetting the solution up and down. The tubes were transferred immediately to a centrifuge at room temperature for a short spin at 200 x g for two minutes. The pellets were washed with HBSS four times before being gently resuspended in 10.0 ml of DMEM (low glucose) medium. The islet suspension was poured into a 15 mm x 100 mm polystyrene petri dish, where the islets were hand-picked under the dissecting microscope, and transferred into fresh media. The islets were incubated at 37 °C with 5.0 % CO_2_ for 48 hours before being used for experiments.

The intragranular pH measurements in β-cells inside isolated islets using DND-160 followed the same procedures we used for cultured cells. After being washed with HBSS, the cells were stained with DND-160 and washed for imaging under a Zeiss LSM-800 confocal microscope. Pairs of images were obtained in tandem together with the DIC image. To avoid potential problems from uneven DND-160 staining, the focal level was limited to the three layers of β-cells right underneath the islet surface. Data analysis followed what was described for INS-1 and PC-12 cells.

### 8. Preparation of monolayer cultures of mouse pancreatic β-cells

Monolayer cultures of primary mouse β-cells were prepared as previously described [75]. Briefly, mouse islets were hand-picked from suspension cultures and washed twice in PBS without Ca^2+^ and Mg^2+^. Dissociation of islets into single cells was performed through continuous, gentle pipetting in 0.30 mL of 0.05% trypsin-EDTA for 3 minutes at 37°C. To prevent additional trypsin digestion, islet monolayer medium was added to a total volume of 15 ml. The islet monolayer medium was prepared from Minimum Essential Medium (MEM) with Gautama, 11 mM glucose, 5.0% FBS, 1.0 mM sodium pyruvate, 10 mM HEPES and 1x B-27 Supplement. Cells were pelleted by centrifugation for five minutes at 350 x g and resuspended in the islet monolayer medium, counted, and seeded in a density of 35,000 cells/cm^2^ on laminin-coated (Gibco; 50 µg/mL in HBSS with Ca^2+^ and Mg^2+^ for 1 hour at 37°C) borosilicate glass coverslips of 12 mm in diameter and 0.17 mm in thickness (Electron Microscopy Sciences) or Mate No. 1.5 35-mm glass-bottomed dishes. Islet cell monolayers are cultured for 3-4 days before experiment.

### 9. *In situ* calibration of ratiometric pH measurements using NPY-ClopHensor

The *in situ* pH calibration was performed as described by Arosio, D. *et al.* [67]. The desired pH was controlled by equilibrating extra- and intra-cellular aqueous phases using nigericin (5.0 μM; a K^+^/H^+^ carrier), carbonyl cyanide m-chlorophenyl hydrazone (CCCP at 5.0 μM; a protonophore), and valinomycin (5.0 μM; a K^+^ ionophore) in the presence of high-K^+^ buffer containing 20 mM HEPES, 0.6 mM MgSO_4_, 38 mM sodium gluconate and 100 mM potassium gluconate. To avoid CO_2_-dependent pH shift during calibrations, cells were maintained in a CO_2_-free atmosphere at 37 °C. The medium pH level was clamped with a proper buffer and the media were changed three times (30 min for each incubation) to ensure stabilization of intracellular pH and intragranular pH. Multichannel fluorescence images (3 – 5 acquisitions, *n* =20–30 independent images) were collected for cells equilibrated at each pH level. Green-to-cyan ratios (R) were expressed as mean ± *s.d*. A polynomial function,

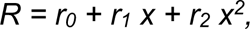

where *x* represents a set buffer pH value, *r_0_, r_1_* and *r_2_* are fitting constants, was used to fit the data and derive an empirical equation for pH readouts.

### 10. Fluorescence imaging and analysis using NPY-ClopHensor

Fluorescence images were collected in a Zeiss LSM710 inverted confocal microscope with an oil-immersion objective (60 ×, NA 1.25) inside a CO_2_ and temperature-controlled chamber. Excitation at 458 nm and 488 nm was achieved by an argon laser. To minimize bleed-through artifacts, emission channels were chosen as follows: Green channel fluorescence I_488 nm_ (488 nm excitation) was detected from 520 nm to 550 nm; cyan channel I_458 nm_ (458 nm excitation) was detected from 500 nm to 550 nm. PMT voltage was kept constant. Laser scanning was performed using 400 Hz line frequency, 512 × 512 (pixel x pixel) format and a pinhole aperture set at 1 Airy disk. Image processing was performed using customized ImageJ (US National Institutes of Health) macros that could be applied to sets of images in a consistent and unbiased way. Mean background fluorescence measured either in non-transfected cells or in empty spaces indicated absence of autofluorescence. The I_488_ / I_458_ ratio was calculated. The equation from non-linear regression fitting of the calibration datapoints were used to calculate pH value of each readout from the granules.

### 11. Analyses of induced granule biogenesis by CHGBΔMIF in CHO and HEK293T cells

The CHO cells were seeded a day (day 0) before transfection at ∼ 30% confluency in glass-bottomed petri dishes. On day 1, cells were transfected with plasmids carrying cDNAs of wild-type CHGB or CHGBΔMIF. Imaging was done 48 hours after transfection. Immediately before imaging, cells were washed with a physiological medium (10 mM HEPES, 120 mM NaCl, 4.8 mM KCl, 2.5 mM CaCl_2_, 1.2 mM MgCl_2_, 24 mM NaHCO_3,_ 3.0 mM glucose, pH 7.4; or 10 mM HEPES, 138 mM NaCl, 4.8 mM KCl, 2.5 mM CaCl_2_, 1.2 mM MgCl_2,_ 3.0 mM glucose and 0.1% BSA, pH 7.4) and then incubated with 1.0 μM DND-160 for 5 min at 37 °C. Excess dye was washed away with the physiological medium and images of the cells were captured under a LSM 780 (or LSM800) upright confocal microscope using an Apochromat 63x/1.40 Oil DIC objective. Images were collected using 405 nm excitation and emissions at 510 nm (yellow) and 485 nm (blue) with 20 nm apertures. ClopHensor was excited at 488 nm and images were captured using emission at 525 ±10 nm.

### 12. Statistics and reproducibility

A strong form of the large number theorem is applicable to the analysis of a large number (> 50) of individual granules randomly selected from blindly chosen cells in every one of the repeated experiments. When hundreds of granules were analyzed, the mean of the observations is close to the expected true values. All experiments were repeated at least 3-4 times independently to assess reproducibility. For statistical comparison of any two chosen datasets, unless separately stated, we applied the standard two-tailed Student *t*-test without the assumption of same variances.

## Abbreviations

CHG: chromogranin
CHGB: chromogranin B
ISG: immature secretory granule
DCSG: dense-core secretory granule
TGN: *trans*-Golgi network
CLC: chloride channel / transporter
RRG: readily releasable granule
MIF: membrane-interacting fragment
CHGB-MIF: CHGB membrane-interacting fragment
CHGBΔMIF: CHGB with its MIF deleted

## ACKNOWLEDGEMENTS

We are grateful to Dr. Barbara Ehrlich (Yale University) for providing the constructs for mouse CHGB, to Dr. Herbert Y. Gaisano (University of Toronto) for the syncollin-pHluorin construct, to Dr. Kuixing Zhang (UC San Diego) for sharing his techniques for releasing granular contents from endocrine cells, to Drs. Ilya Bezprozvanny, Kate Phelps, and Jen Liou (UT Southwestern Medical Center) for their assistance and advice on the intragranular pH measurements using ratiometric dyes, to Dr. Wen-Hong Li for sharing the murine INS-1 cells and the insulin detection methods, to Dr. Jerry Shay (UT Southwestern Medical Center) for providing the Lentiviral construct of CHGB shRNA, and to Dr. Edwards Phelps at the University of Florida for technical guidance on the experiments with β-cell monolayers. We thank Dr. Mahesh Chandak and Ms. Sutonuka Bhar (University of Florida) for their technical assistance.

## Ethics approval

The authors confirm that all experiments with mice were performed by following strictly the rigorous protocols (#201709886) developed by the Jiang lab and approved by the IACUC committee at UF. The mouse colony raised out of frozen sperms and bred by the vendor were first quarantined and then bred in the UF animal care service facility. All mice used for experiments were moved out the facility on the morning of the planned surgery day and were sacrificed on the same day.

## Consent for publication

All authors agree to publish this paper.

## Availability of data and materials

All data are available upon request. We will share the constructs and the information of the animal model with all requesters.

## Author Contributions

Q.-X.J. designed and oversaw the experimental studies, analyzed all results with co-authors and did drafting and revision of the manuscript. G.Y. and Q.X.J. designed and performed all molecular cloning, biochemical studies, and cell-based experiments, and analyzed all acquired data. M.A. and G.Y. performed dissection of mice and isolation of pancreatic islets with C.M.’s advice. D.W.H provided technical guidance and assistance in preparation of monolayers of pancreatic β-cells in the Phelps lab. L.C. helped with the mouse model studies of the results. All authors contributed to data analysis and commented on manuscript.

