## Supplemental Information for "Molecular requirements of chromogranin B for the long-sought anion shunter of regulated secretion"

#### Supplementary Information

**I. Supplementary Figures**
**II. List of reagents**
**III. Supplementary Table S1**
**IV. Supplementary References**

Confidential

### I. Supplementary Figures

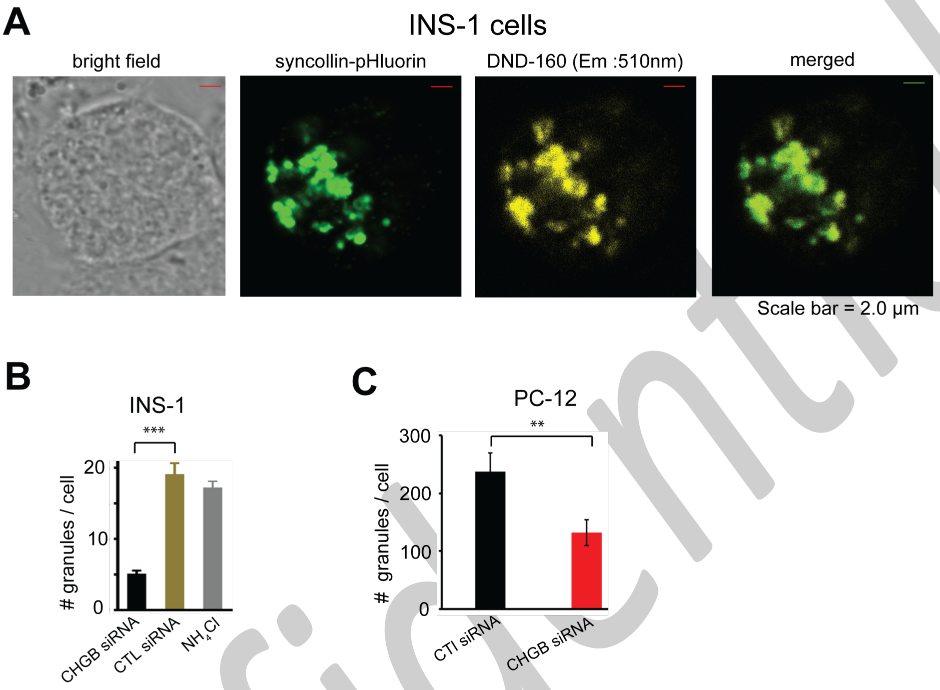


Confidential

**Supplementary Figure 1. DND-160 for staining secretory granules in endocrine cells.**

**(A).** Syncollin-pHluorin, a granule protein fused with a fluorescent tag, was expressed in INS-1 cells to examine if DND-160 stains secretory granules well. Cells were stained with 0.5 $\mu$M DND-160 and imaged. Typical DIC image (1$^{st}$), syncollin-pHluorin (2$^{nd}$) and DND-160 (3$^{rd}$) images of the same cell and merging of the last two (4$^{th}$) are showed. Syncollin and DND-160 overlap very well, suggesting that nearly all DND-160-stained compartments are syncollin-positive secretory granules, mainly because the granules outnumber other acidic compartments like lysosomes in these cells. **(B).** Average numbers of DND-160-stained granules per INS-1 cell section from cells treated differently and imaged under a confocal microscope. Cells transfected with CHGB-targeting or control siRNAs or those treated with 5.0 mM $NH_4Cl$ were analyzed. Granules in ~40 cells were counted for each condition. Error bar: *s.d.* ***: $p < 0.001$. (**C**). Average numbers of SGs per PC-12 cell section transfected with CHGB-specific or control siRNAs. Error bars: *s.d.*, n = 35. **: $p < 0.01$.

Confidential

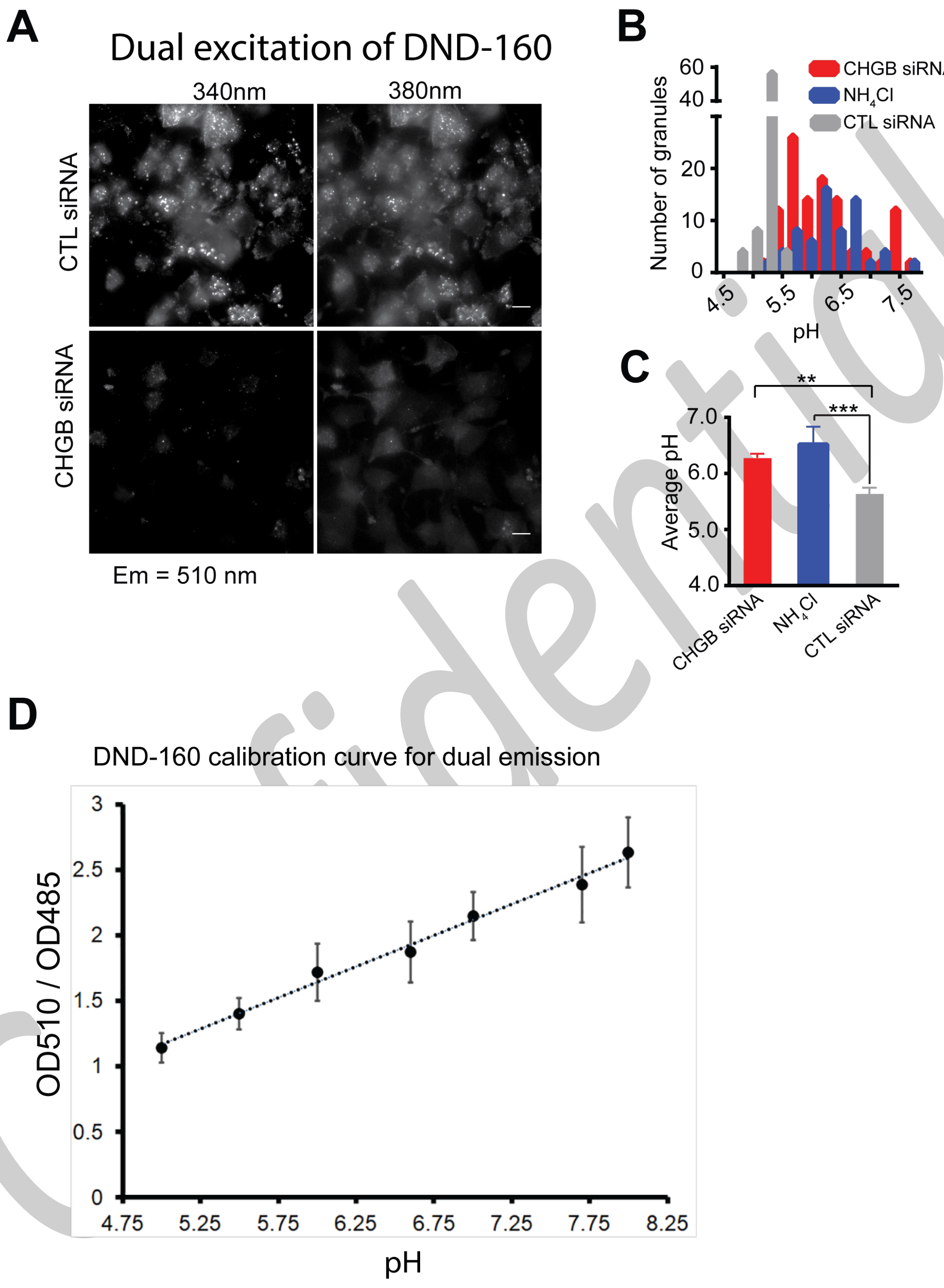


Confidential

**Supplementary Figure 2. Dual excitation measurements of DND-160-stained granules in INS-1 cells.**

(**A**). Cells were treated with control (CTL; top row) and CHGB-specific siRNAs (bottom row), respectively and imaged at 510 nm after being excited in sequence at 340 and 380 nm. (**B**). Histograms of granular pH values from ratiometric measurements of dual-excitation images of cells transfected with CTL siRNA (gray bars), CHGB-targeting siRNA (red bars) and those treated with 5.0 mM $NH_4Cl$ (blue bars). (**C**). Average intragranular pH values measured from cells differentially treated in **B**. Errors: *SEM*; n=3. Results from the dual-excitation experiments agreed with what was reported before [1]. **(D**). A typical calibration curve of DND-160 for a dual emission experiment. (Error bars: *s.d.*; n = 4). The linear regression fitting function was used for reading out pH for each specific measurement. The calibration was done for every imaging system.

Confidential

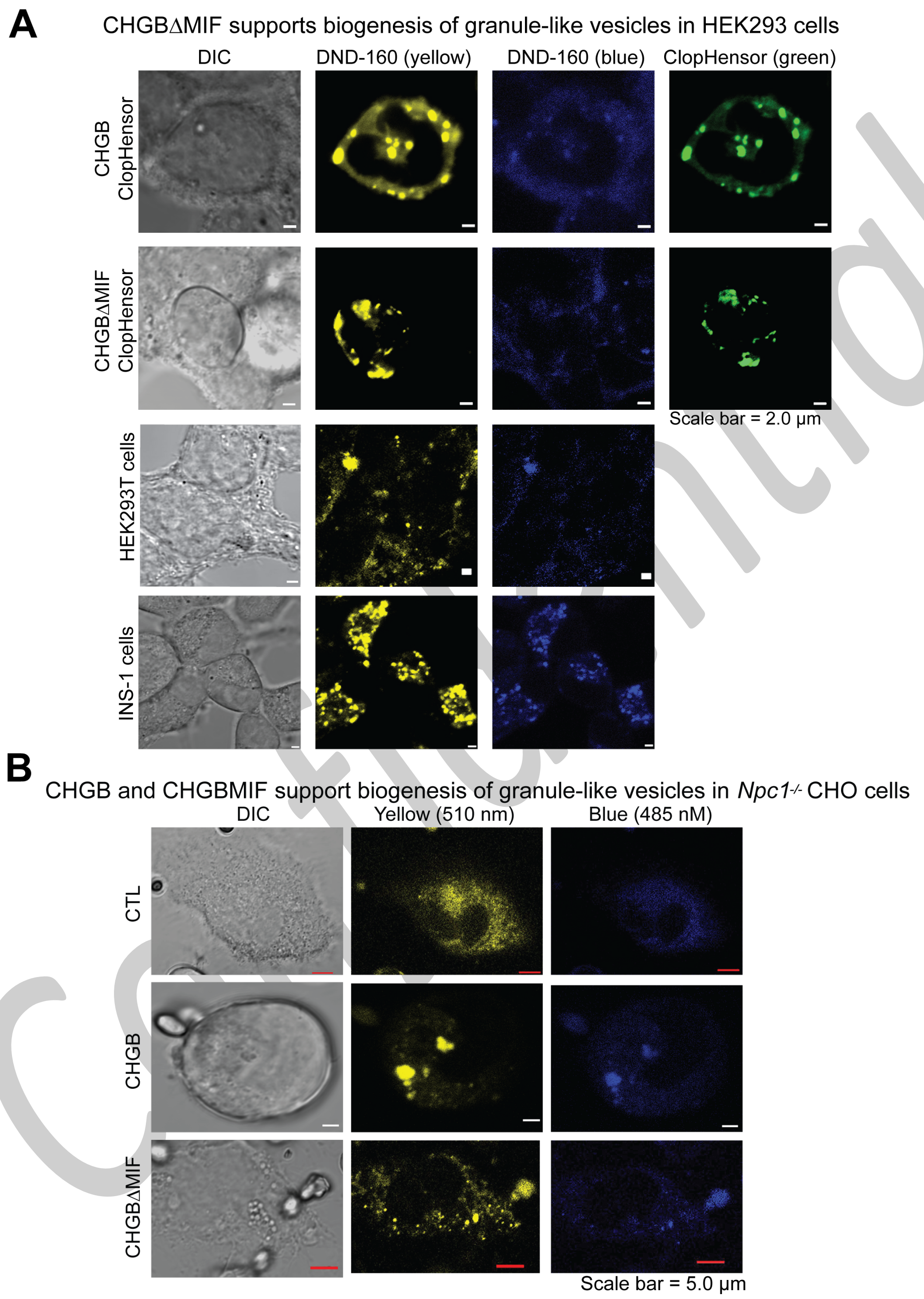
A
CHGBΔMIF supports biogenesis of granule-like vesicles in HEK293 cells
DIC
DND-160 (yellow)
DND-160 (blue)
ClopHensor (green)
CHGB ClopHensor
CHGBΔMIF ClopHensor
HEK293T cells
INS-1 cells
Scale bar = 2.0 µm
B
CHGB and CHGBMIF support biogenesis of granule-like vesicles in Npc1-/- CHO cells
DIC
Yellow (510 nm)
Blue (485 nM)
CTL
CHGB
CHGBΔMIF
Scale bar = 5.0 µm

**Supplementary Figure 3. CHGB and CHGBΔMIF induce biogenesis of secretory granule-liked vesicles (SGLVs) in cultured fibroblast cells.** (**A**). HEK293T cells overexpressing CHGB-ClopHensor or CHGBΔMIF-ClopHensor fusion proteins showed SGLVs. ClopHensor was attached to the C-terminal end of CHGB or CHGBΔMIF. Besides ClopHensor signals, cells were acutely labeled with DND-160 for imaging in a Zeiss LSM800. **Row 1**: HEK293T cells over-expressing CHGB-ClopHensor. DND-160 and ClopHensor signals overlap at every spot. **Row 2**: HEK293T cells overexpressing CHGBΔMIF-ClopHensor also show nearly perfect agreement between DND-160 and ClopHensor signals. **Row 3**: HEK293T cells labeled with DND-160 as a negative control. The background staining appears on the cell membrane, not inside the cells. **Row 4**: INS-1 cells labeled with DND-160 as a positive control, showing patterns of granules very different from that of the control cells in Row 3. (**B**). Images of $Npc1^{-/-}$ CHO cells overexpressing CHGB or CHGBΔMIF. Cells were labeled with DND-160 for dual-emission imaging. **Row 1 (CTL):** control cells. **Row 2 (CHGB)**: cells overexpressing CHGB. **Row 3 (CHGBΔMIF)**: cells overexpressing CHGBΔMIF. The bigger bright puncta are the SGLVs containing CHGB or CHGBΔMIF.

Confidential

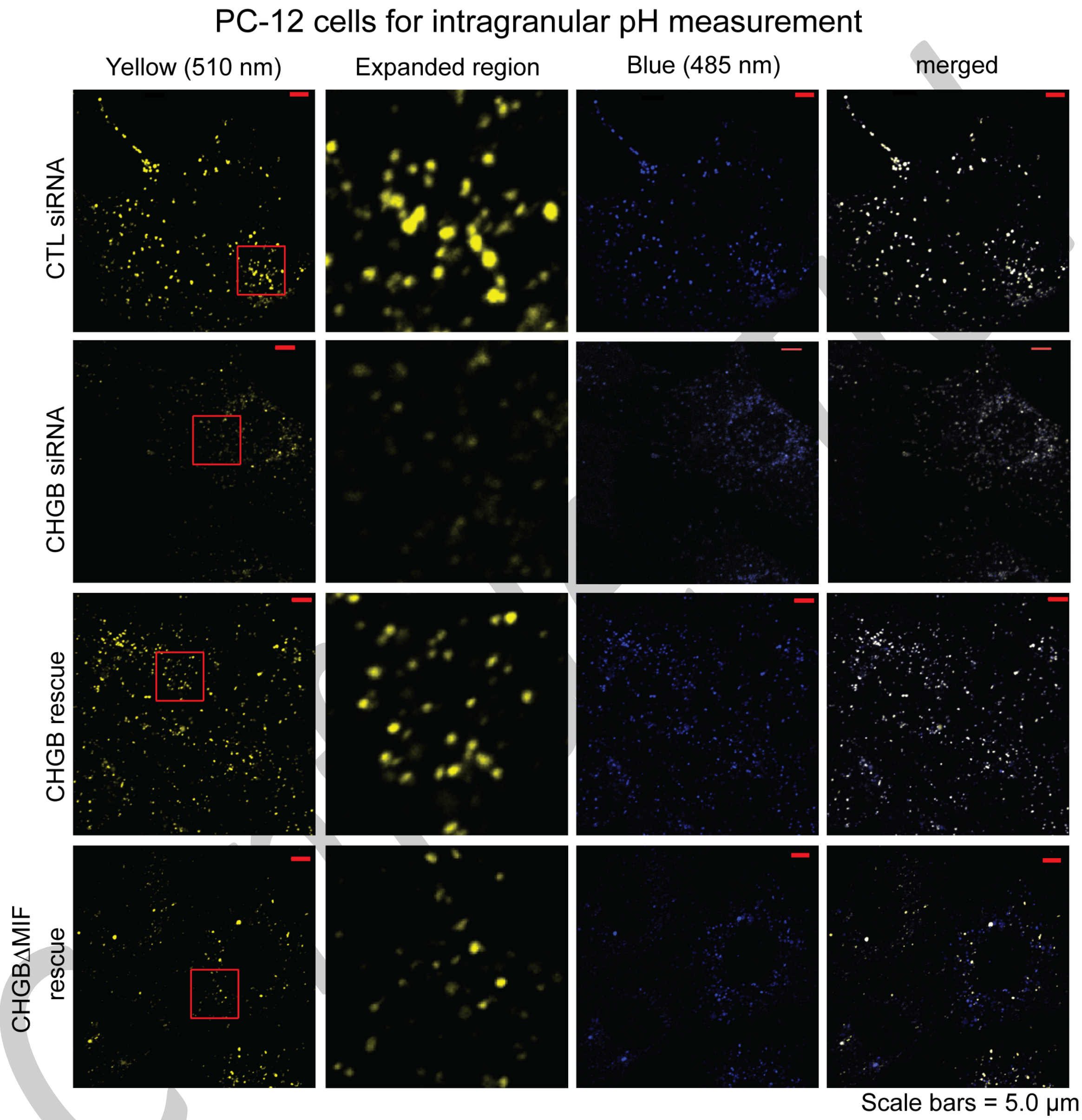
PC-12 cells for intragranular pH measurement
Yellow (510 nm)
Expanded region
Blue (485 nm)
merged
CTL siRNA
CHGB siRNA
CHGB rescue
CHGBΔMIF rescue
Scale bars = 5.0 μm

**Supplementary Figure 4. Typical images of DND-160-stained PC-12 cells in four different conditions for ratiometric measurements of intragranular pH**.

After being transfected with CTL or CHGB-targeting siRNAs, the cells were incubated for two days before some of the CHGB-knockdown cells were transfected to overexpress CHGB or CHGBΔMIF. All cells were fed with fresh media for two more days before DND-160 imaging. PC-12 cells transfected with control or CHGB-specific siRNAs are in the $1^{st}$ and $2^{nd}$ rows, respectively. Rows 3 & 4 show CHGB-knockdown cells that were rescued by over-expressing CHGB (3rd) or CHGBΔMIF ($4^{th}$). Excitation: 410 nm; emission: 485 nm and 510 nm. A small area marked by a red window in each 510 nm image (yellow) was expanded to show granules (column 2). Measurements at 510-nm in rows 2 & 4 were weaker due to pH change.

Confidential

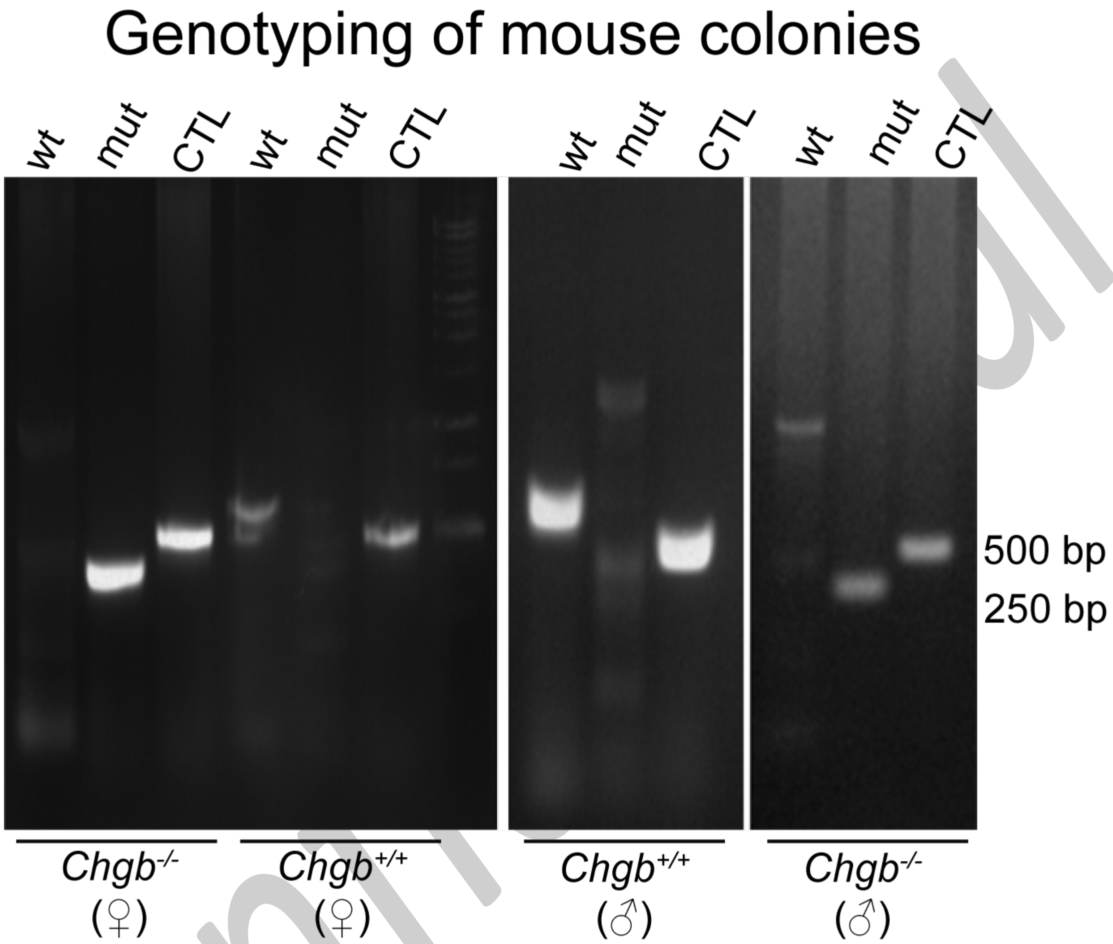
Genotyping of mouse colonies
wt
mut
CTL
wt
mut
CTL
wt
mut
CTL
wt
mut
CTL
500 bp
250 bp
$Chgb^{-/-}$ (♀)
$Chgb^{+/+}$ (♀)
$Chgb^{+/+}$ (♂)
$Chgb^{-/-}$ (♂)

**Supplementary Figure 5. Genotyping data from *Chgb*$^{-/-}$ and control mice**

Mouse colonies used for experiments were examined by PCR of tail nips which were sampled at 4-weeks of age. The wild-type and knockout alleles give rise to different patterns of PCR fragments after electrophoresis. Typical data for a pair of male and female littermates of homozygous mice and a pair of wild-type mice are presented.

Confidential

#### II. List of reagents

| REAGENT or RESOURCE | SOURCE | IDENTIFIER |
|---|---|---|
| **Antibodies** | | |
| anti-chromogranin B antibody (Santa Cruz) | Goat | sc-1489 |
| anti-chromogranin B antibody (Santa Cruz) | Rabbit | SC-20135 |
| anti-His antibody (Sigma) | Mouse | 27471001 (lot#9535913) |
| anti-CLC-3 antibody | Rabbit | ab28736 (GR383528-I |
| anti-chromogranin B | Rabbit | PA1-10839 (TI2638713) |
| Anti-chromogranin A | rabbit | H300, new sc-13090 |
| anti-ATP6V0A2 | Rabbit | ab96803 (GR9499-19) |
| **Bacterial and Virus Strains** | | |
| *E. coli* | available in the lab | K-12 (XL1 blue) |
| Baculovirus | Invitrogen | Bac-to-Bac expression system |
| DH10Bac | Invitrogen | Cat #10361012 |
| **Biological Samples** | | |
| Mouse islets | Isolated in the lab | |
| **Experimental Models: Cell Lines** | | |
| INS-1 1832/13 cell line | Wen-Hong Li's lab at UT Southwestern | INS-1 |
| PC-12 cell line | Jerry Shay's lab at UT Southwestern | PC12 |
| **Experimental Models: Organisms/Strains** | | |
| Mouse | InfraFrontier GmbH in Germany | **C57BL/6NTac-Chgb**$^{tm1(EGFP/cre/ERT2)Wtsi}$**/WtsiIeg** |
| **Recombinant DNA** | | |
| pPCDNA3.0-CHGB | constructed in the lab | CHGB |
| pPcDNA3.0-CHGB ΔMIF | constructed in the lab | CHGBΔMIF |
| pcDNA3-NPY-ClopHensor | Addgene.com | Plasmid #25939 |
| pcDNA3-ClopHensor | Addgene.com | Plasmid #25938 |
| pfastBac1 | Invitrogen | Cat # 10360014 |
| **Software and Algorithms** | | |
| ImageJ | NIH | https://imagej.nih.gov/ij/ |

Confidential

**III. Supplementary Table S1.** Missense polymorphism in *Chgb loci* associated with human Type 2 diabetes (from published database [2]).

| "Variant ID" | dbSNP ID | Major allele | Minor allele | Predicted impact | Residue change from the seq. in NCBI | p-Value | Effect | MAF |
|---|---|---|---|---|---|---|---|---|
| 20_5843952_G_A | rs41282138 | G | A | missense | T154Q | 0.013 | 0.464 | 0.011 |
| 20_5903848_C_G | rs236152 | C | G | missense | A353G | 0.019 | 1.16 | 0.38 |
| 20_5904028_C_T | rs742710 | C | T | missense | P413L | 0.03 | 1.08 | 0.15 |
| 20_5904300_A_G | rs140131066 | A | G | missense | K504E | 0.032 | 2.96 | 0.0085 |
| 20_5903932_G_T | rs142841879 | G | T | missense | W381L | 0.034 | 0.624 | 0.016 |
| 20_5903067_T_A | rs6085324 | T | A | missense | S93T | 0.035 | 0.861 | 0.28 |
| 20_5903141_G_C | rs236150 | G | C | missense | K117N | 0.038 | 1.06 | 0.025 |
| 20_5904040_G_A | rs742711 | G | A | missense | R417H | 0.041 | 0.834 | 0.39 |
| 20_5843918_C_T | rs74824950 | C | T | missense | P143C | 0.043 | 10.9 | 0.00084 |
| 20_5903323_G_A | rs910122 | G | A | missense | R178Q | 0.056 | 0.882 | 0.38 |
| 20_5903388_A_C | rs881118 | A | C | missense | N200H | 0.057 | 1.16 | 0.17 |
| 20_5904289_G_A | rs74621755 | G | A | missense | R500K | 0.072 | 1.85 | 0.022 |
| 20_5903485_G_A | rs6139873 | G | A | missense | R232Q | 0.074 | 0.64 | 0.031 |
| 20_5843672_A_G | rs114118253 | A | G | missense | L61A | 0.087 | 9.23 | 0.0012 |
| 20_5903517_A_G | rs236151 | A | G | missense | T243A* | 0.096 | 1.03 | 0.87 |

Notes:

T154 is in NCBI database, not R154.
P143 is in NCBI database, not R143.
L61 is in NCBI database, not T61.
A243 is already in the NCBI database, not T243.
**: CHGB peptides:
GAWK (CHGB420-493):
CCB (CHGB597-653); and
secretolytin (CHGB614-626).
***: CHGB-MIF is CHGB440-594.

Confidential

Confidential
